# Sex differences in bile acid homeostasis and excretion underlie the disparity in liver cancer incidence between males and females

**DOI:** 10.1101/2020.06.25.172635

**Authors:** Megan E. Patton, Sherwin Kelekar, Lauren J. Taylor, Angela E. Dean, Qianying Zuo, Rhishikesh N Thakare, Sung Hwan Lee, Emily Gentry, Morgan Panitchpakdi, Pieter Dorrestein, Yazen Alnouti, Zeynep Madak-Erdogan, Ju-Seog Lee, Milton J. Finegold, Sayeepriyadarshini Anakk

**Author notes:** Department of Chemistry, Virginia Tech, Blacksburg, VA 24061.

## Abstract

Hepatocellular carcinoma (HCC), the most common liver cancer, exhibits a higher incidence in males. Here, we report that mice lacking the bile acid regulators, Farnesoid X Receptor (FXR) and Small Heterodimer Partner (SHP), recapitulate the sex difference in liver cancer risk. Since few therapeutic options are available, we focused on understanding the intrinsic protection afforded to female livers. Transcriptomic analysis in control and FXR and SHP double knockout livers identified female-specific changes in metabolism, including amino acids, lipids, and steroids. We examined if the obtained transcriptomic signatures correlate with the survival outcomes for HCC patients to assess the translational potential of this murine HCC model. Gene signatures that are unique to the knockout females correspond with low-grade tumors and better survival. Ovariectomy blunts the metabolic changes in female livers and promotes tumorigenesis that, intriguingly, coincides with increases in serum bile acid (BA) levels. Despite similar genetics, we found higher serum BA concentrations in males, whereas female knockout mice excreted more BAs. Decreasing enterohepatic BA recirculation using cholestyramine, an FDA-approved resin, dramatically reduced the liver cancer burden in male mice. Overall, we reveal that sex-specific BA metabolism leading to lower circulating BA concentration protects female livers from developing cancer. Thus, targeting BA excretion may be a promising therapeutic strategy against HCC.

**Significance:** We show that female-specific gene profiles identified in *Fxr*^-/-^, *Shp^-/-^* double knockout (DKO) mice correlate with better outcomes for HCC patients and uncover sex differences in circulation and excretion of bile acid. Overall, we demonstrate that increasing (ovariectomy or chemical injury) or decreasing (pharmacologically with FDA-approved resin) serum bile acids, not hepatic bile acids, promoted or alleviated liver cancer burden.

## Introduction

Liver cancer, a leading cause of cancer-related death, has diverse etiologies and displays sex-difference with reduced risk in females compared to males [1–5]. Since current therapies for liver cancer fall short, we posit that understanding molecular mechanisms functioning in the female livers will reveal new therapeutic targets. Earlier studies have reported the role of sex hormones [6–9], transcription factors FoxA1/A2 [10], and cytokine Il6 signaling [11] in regulating the sex difference in hepatocellular carcinoma (HCC), but the role of metabolic pathways remains poorly understood.

Rewiring of cellular metabolism enables the tumor cells to maintain viability and grow disproportionately [12]. We previously showed that the combined deletion of nuclear receptors, Farnesoid X Receptor (FXR), and Small Heterodimer Partner (SHP) resulted in spontaneous liver cancer in the year-old male mice [13]. In this study, we report that, unlike the males, female *Fxr*^-/-^, *Shp^-/-^*double knockout (DKO) mice exhibit protection against tumorigenesis and thus mimic the sexual dimorphism in liver cancer incidence observed in clinics. Although 15-month-old individual *Fxr* knockout and individual *Shp* knockout mice were previously shown to develop liver cancer, unlike the DKO mice, their incidence does not show 100 percent penetrance nor sex differences [14–16].

Mutations and reduction in *Fxr,* and *Shp* transcript levels have been noted in cholestasis (reduced bile flow and subsequent increase in hepatic and serum bile acids (BA)), fatty liver disease, and liver cancer [17–23]. Moreover, individuals with chronic cholestasis exhibit an increased risk for HCC [24–26]. Typically, BA levels are tightly controlled via receptor signaling, including FXR and SHP [27–31]. Consistently, combined loss of *Fxr* and *Shp* in mice results in juvenile onset cholestasis that progresses to HCC [32]. We and others have shown that excessive accumulation and dysregulation of BA homeostasis are directly linked with liver cancer risk [13, 26, 33–35]. However, whether BAs are contributing factors to the sex differences seen in HCC prevalence has not been evaluated.

Therefore, we performed transcriptomic analysis to identify distinct gene profiles from both sexes of control and DKO mice. Then, using five separate human clinical HCC cohorts, we tested the clinical utility of the identified gene signatures from our mouse model. Next, we investigated the role of endogenous estrogen signaling in the DKO mice by performing ovariectomy. We measured hepatic, serum, urine, and fecal BAs from male and female mice to understand their homeostasis. Finally, we manipulated the circulating BA levels in the DKO mice either with a chemical challenge or BA binding resins and examined its consequence on hepatic tumorigenesis. Overall, our data uncover that the differential BA homeostasis between the two sexes can orchestrate the observed gender differences in HCC burden in clinics.

## Results

### Fxr^-/-^, Shp^-/-^ double knockout (DKO) mice phenocopy clinical features of HCC

Here, we report that DKO mice exhibit the sexually dimorphic incidence of HCC observed in the clinic. Despite the loss of BA homeostatic machinery, one-year-old DKO female mice did not develop liver tumors but showed modest fat accumulation and mild fibrosis. On the contrary, DKO male livers revealed HCC and well-defined adenomas, robust steatosis, and fibrosis (Fig. 1A-F). Even at six months of age, female livers were smaller and displayed reduced hepatic fibrosis compared to males (Supplementary Fig. 1). Difference in tumor burden was reflected in the gross liver to body weight ratio, which was significantly higher in DKO males than DKO females (Fig. 1G). Serum ALT and AST were elevated in the DKO animals compared to WT (Fig. 1H) and correlated with the cholestatic phenotype of the mice. More importantly, liver cancer patients showed a reduction in *Fxr* and *Shp* transcript expression (Fig. 1I).

**Figure 1:**
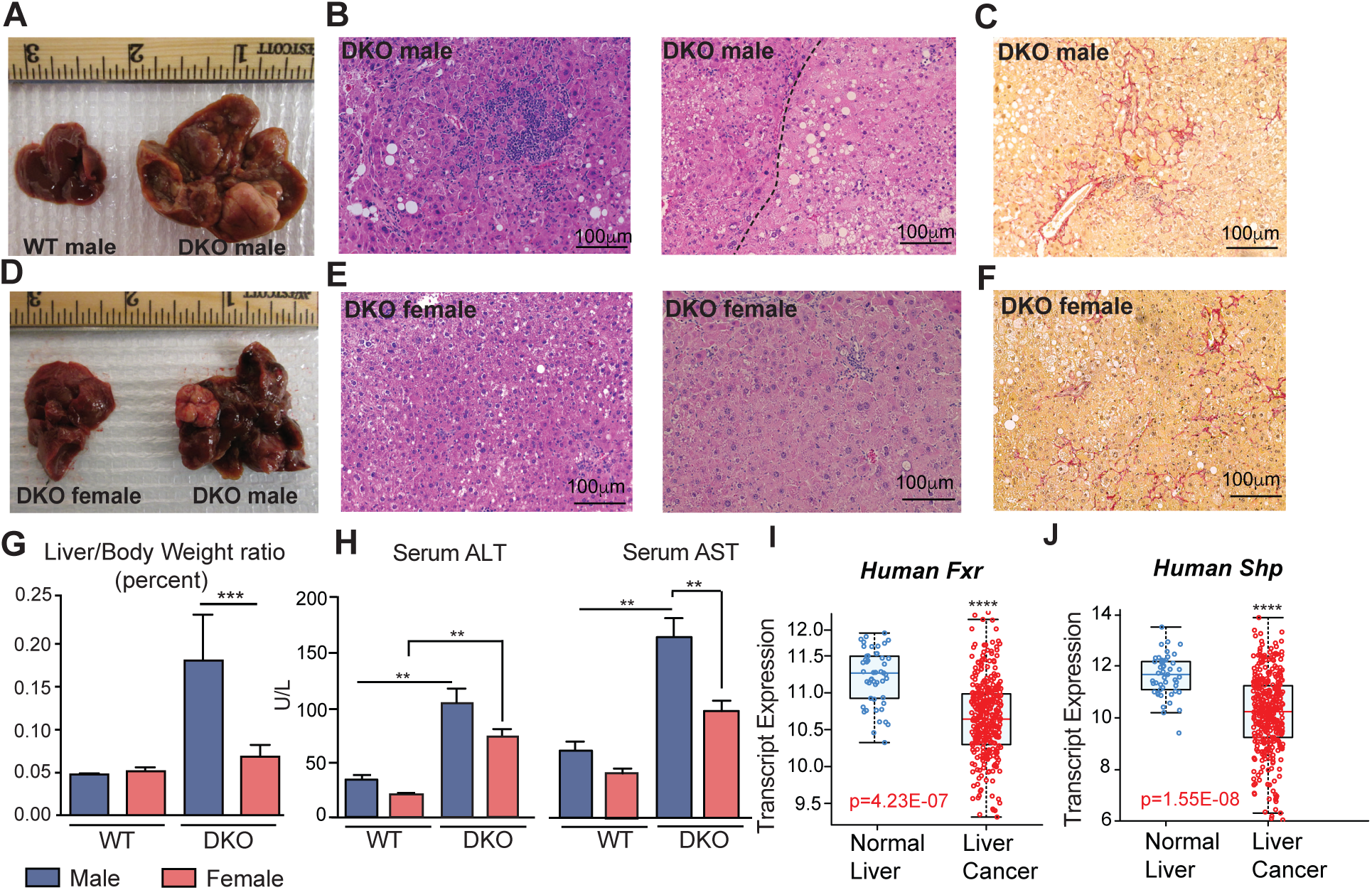
*Fxr/Shp* double knockout (DKO) mouse model recapitulates sexdifference observed in HCC incidence. (A-E) One-year-old DKO male mice developed hepatocellular carcinoma, which was not observed in age-matched wild type (WT) and DKO female mice. (B and E) Representative H&E stained liver sections from a (B) DKO male and (E) female. Inflammation and injury are evident at 1 year, and the dotted line (B) separates the HCC with large nuclei on lower right. Sirius red staining shows increased collagen in a perisinusoidal distribution, which is greater in the DKO males (C and F). The liver-to-body weight ratio was significantly higher in DKO males (G). Compared to WT animals, serum markers of liver injury, (H) AST and ALT were higher in DKO mice. (I-J) Analysis of five different HCC clinical cohorts (n=1000) patients reveals a reduction in *Fxr* and *Shp* transcript levels in patients with liver tumors. n=5-10 mice /group; mean ± SEM; *p<0.01, **p<0.001 compared to genotype or gender controls. One-way ANOVA with Bonferroni post hoc analysis was performed.

### Sex-specific metabolic programs regulate liver tumorigenesis

To identify transcriptional mechanisms that can contribute towards the sex differences in the incidence of hepatic tumorigenesis, we analyzed one-year-old male and female livers. DKO males and females displayed striking differences in hepatic gene expression profile (Fig. 2, and GEO GSE151524), with DKO males showing enrichment of endoplasmic reticulum stress, unfolded protein response and immune function (Fig. 2A-B). Additionally, network analysis with ClueGO [36] revealed interactions between drug metabolism, inflammation, ERK signaling, and steroid metabolism in DKO males (Fig. 2C). On the contrary, DKO females displayed pathway enrichment of steroid metabolism and clustering of lipid, glucose, amino acid, and steroid metabolism, along with increased sulfotransferase activity (Fig. 2D). Next, we parsed the sex-specific upregulated gene sets to identify unique transcription factor motifs. Overrepresented motifs in DKO males, including AR, FOXA1, FOXA2, NRF2, and PPARγ [10, 37–40], correlated with tumor-promoting functions (Supplementary Table 2). In contrast, in DKO females, FOXO1, E2F, and ERα (Supplementary Table 3) were dominant motifs and are associated with regulating metabolic function during liver carcinogenesis [41].

**Figure 2:**
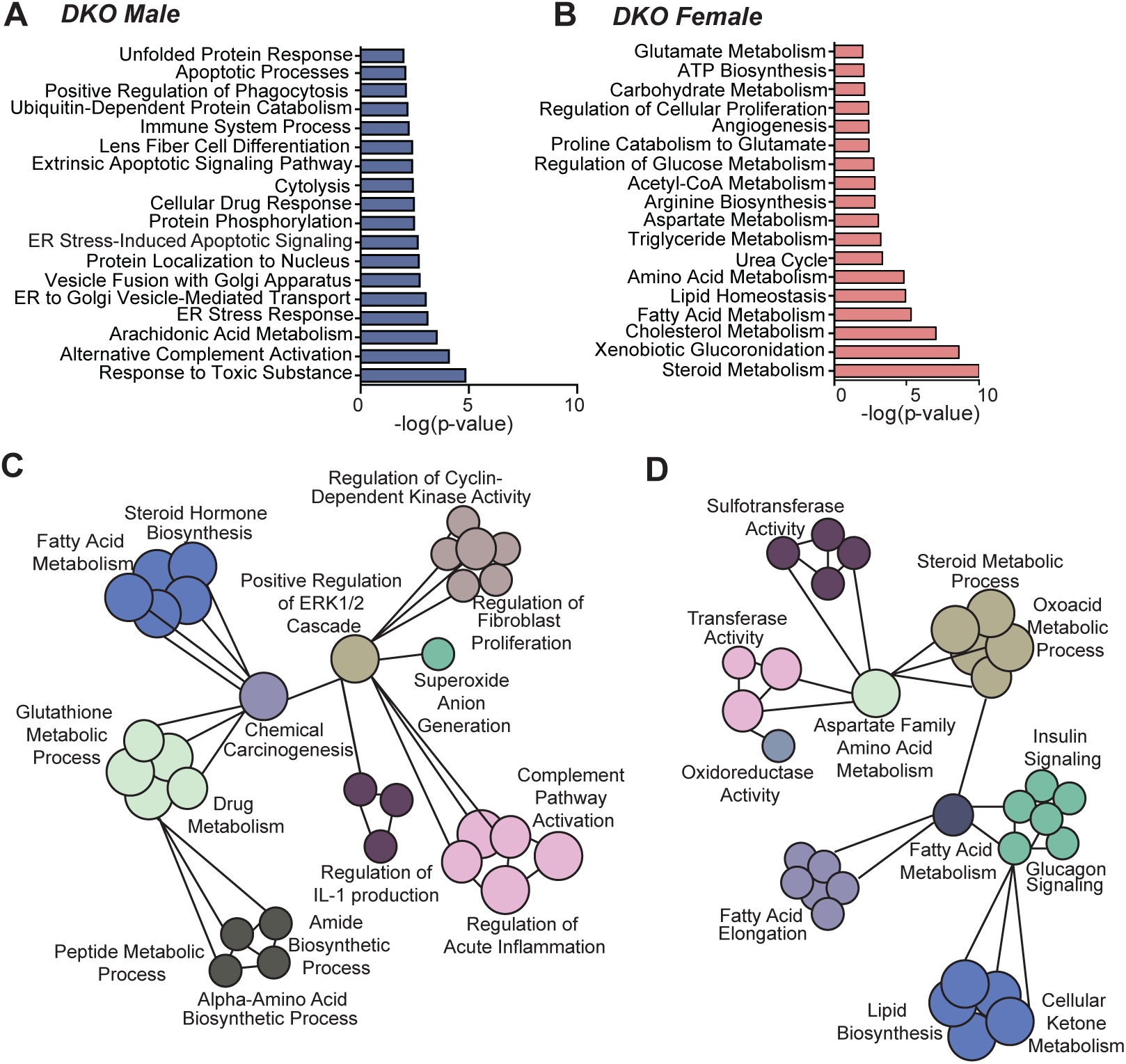
Transcriptome analysis reveals striking sex differences in hepatic metabolism. Microarray was performed on liver tissue from WT and DKO mice of both sexes (*n* = 6/group). (A-B) GO categories were determined using genes with >1.3 fold change in expression between DKO males and females. Enrichment of overlapping GO categories between males and females was determined by comparing – log p-values for each term. (C) GO categories unique to the set of genes upregulated >1.3 fold in DKO males and (D) DKO females.

### The transcriptomic signature of the DKO mice correlates with poor overall survival in the clinical datasets

To investigate the clinical relevance of the DKO mouse model, we analyzed the WT and DKO murine transcriptomic signatures in a sex-specific manner and compared these to five separate clinical HCC datasets (Supplementary Table 4). The patient data were sorted based on similarity to one-year-old DKO gene signatures using class prediction (Supplementary Fig. 2-3). Computational prediction scores (BCCP: 1 represents complete match and 0 represents no match) using the patient samples revealed that the DKO male signature matched with the later stages (≥2) of liver cancer, whereas the DKO female-specific signature matched well with earlier tumor stages (CLIP score 0 or TNM stage 1) (Supplementary Fig. 3).

Of the 1100 patient data, we found approximately (∼45%) showed a transcriptomic signature similar to that of either DKO male or DKO female, which corresponded to lower overall survival (OS), but not recurrence-free survival (RFS). WT gene signatures were used as controls (Fig. 3A-B). Although DKO female mice do not develop liver cancer, it is pertinent to note that they lack *Fxr* and *Shp* expression, display chronic cholestasis similar to their male counterparts, and hence the global gene changes associate with poor OS. On the contrary, when we focused on the gene signature that was distinctly changed only in the DKO female livers, not the DKO males, we found that patients (∼54.71%) who displayed this subset of gene signature had better OS as well as RFS (Fig. 3C). These findings underscore a high potential for clinical translation of data generated from the DKO mouse model. Moreover, by focusing on specific transcript changes in the DKO female livers, we uncovered a subset of metabolic genes that correspond to better survival and might be responsible for their protection against cancer.

**Figure 3:**
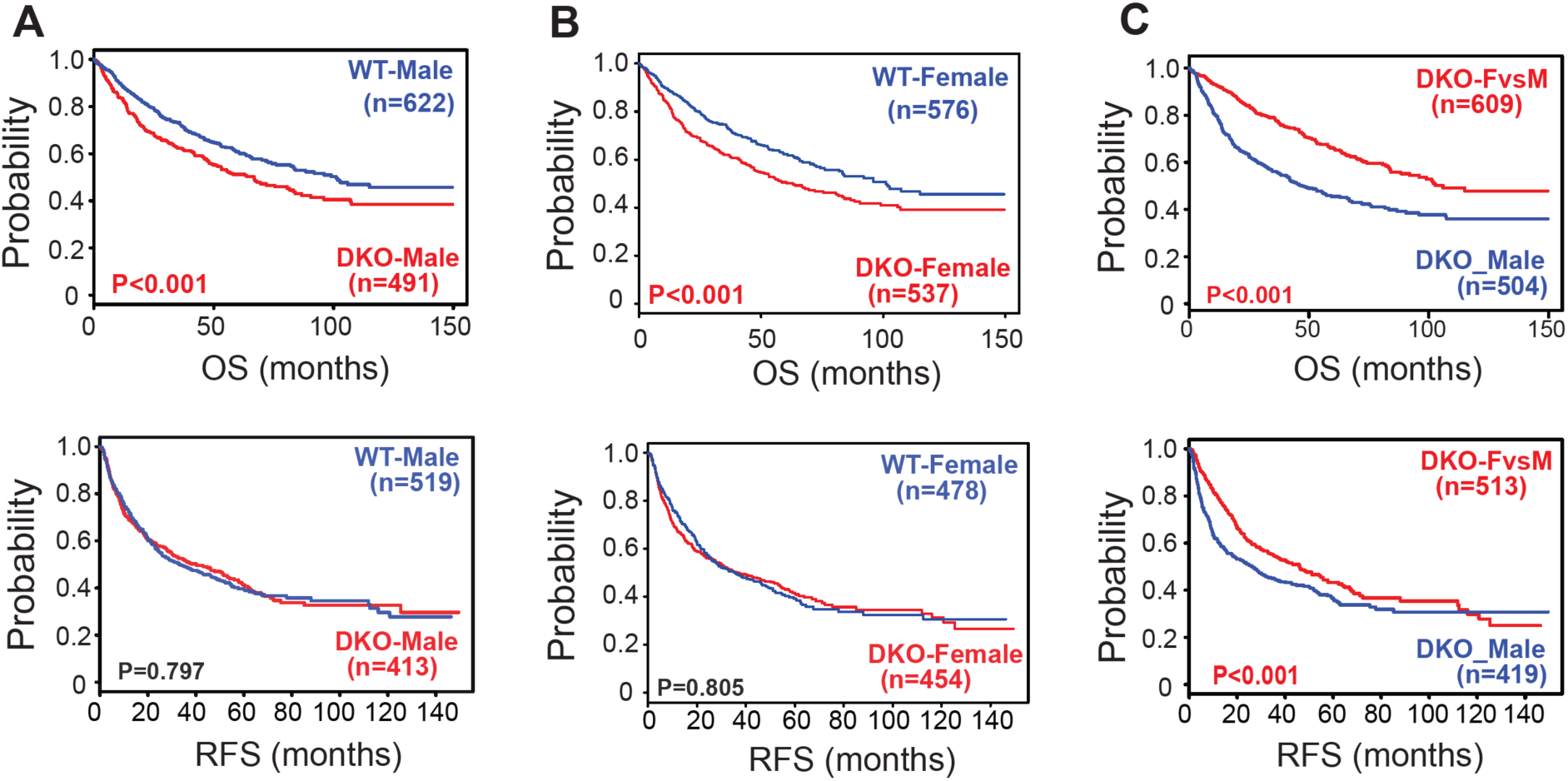
Gene signatures obtained from the DKO mouse model correlate well with the clinical outcomes of HCC patient. The survival probability based on WT and DKO transcriptome changes was evaluated using five different HCC clinical cohorts. (A-C) Analysis of OS (Overall Survival) and RFS (Recurrence Free Survival) in patients using the gene signatures representative of either male WT or DKO mice (A), female WT or DKO mice (B), and unique changes observed in female DKO mice (C).

We initially examined the pathways pertaining to amino acid metabolism and ureagenesis since individuals with mutations in the urea cycle disorder have an increased risk of developing liver cancer [42–45]. Consistent with this, analysis of the TGCA-LIHC clinical dataset revealed a broad downregulation of genes encoding the entire urea cycle, including carbamoyl phosphate synthetase (*Cps1*), ornithine transcarbamylase (*Otc*), argininosuccinate synthetase (*Ass1*), argininosuccinate lyase (*Asl*), and arginase (*Arg1*) in both sexes upon liver tumorigenesis (Supplementary Figure 4). In contrast, these genes were all upregulated in DKO female livers (Supplementary Fig. 5A), which correlated well with the protection afforded to the DKO female livers as loss-of-function mutations in these genes are linked to HCC [45–47]. Additionally, our analysis showed that patients with increased expression of urea cycle genes (DKO-UreaCycle) exhibited a better clinical outcome (Supplementary Fig. 5B).

### Estrogen signaling controls amino acid and bile acid metabolism in the liver

Since estrogen signaling was previously shown to regulate amino acid metabolism [48], we examined its role in controlling the expression of urea cycle genes in the DKO female livers. To do this, we ovariectomized (OVX) DKO mice and found that, indeed the hepatic expression of all these genes, *Cps1*, *Asl1, Ass, Otc* and *Arg1* were significantly blunted in the absence of endogenous estrogen signal (Supplementary Fig. 5C). But when we measured the urea cycle metabolites, we did not find any significant change in the intermediate nor urea production except for a decrease in ornithine levels (Supplementary Fig. 5D), in DKO females compared to the DKO males. We reason that static measurements may not reflect the flux into the urea cycle.

Besides amino acid metabolism, estrogen signaling has been shown to affect BA homeostasis and cause cholestasis [49–51]. So, we anticipated that OVX would lower BA levels in DKO female mice. Instead, we found that OVX led to liver cancer development in otherwise resistant year old DKO female mice (Fig. 4A-B). Moreover, their serum BA levels doubled (Fig. 4C), consistent with the tumorigenic role of BAs. Also, analysis of TCGA-LIHC clinical data revealed significant downregulation of *Erα* gene expression in liver tumors (Supplementary Figure 6A). In addition, estrogen signaling gene signature obtained from the DKO livers correlated with better overall- and recurrence-free survival (Supplementary Figure 6B). Importantly, these results corroborated well with clinical observations that post-menopausal women exhibit higher susceptibility to developing HCC, which can be mitigated upon hormone replacement therapy [52, 53].

**Figure 4:**
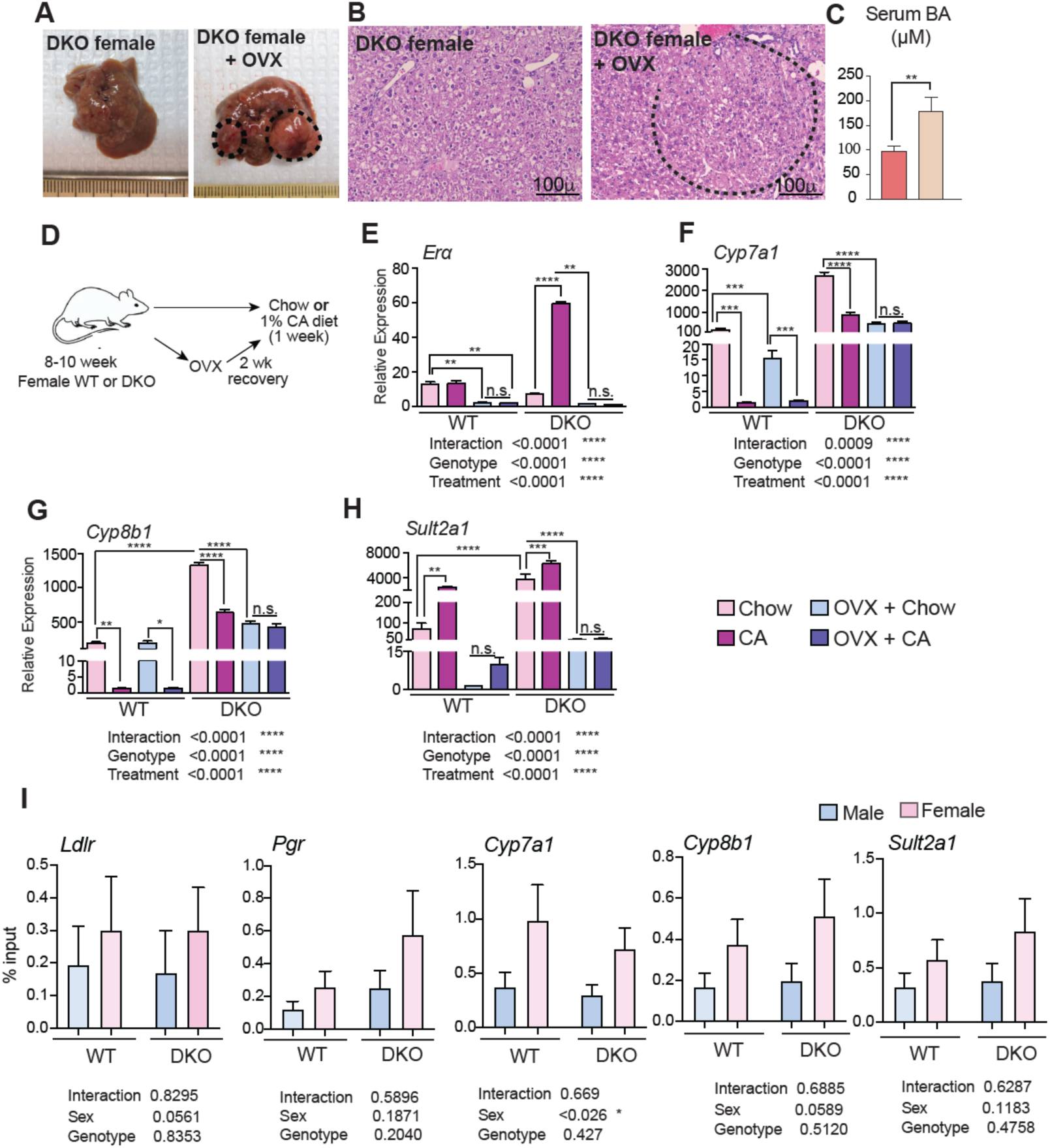
Estrogen signaling protects against liver tumorigenesis and can regulate BA synthesis in DKO female mice. (A-B) Ovariectomized female DKO mice were aged to a year and examined for liver tumorigenesis, where a dotted line demarcates the tumor margin. (C) Serum total bile acid concentrations. (D) Experimental design of chow and 1% cholic acid (CA) diet for 1 week with or without (OVX). (E) Expression of hepatic *Era* was induced *with* CA diet in DKO female mice and reduced in both WT and DKO females following ovariectomy. (F) CA-mediated suppression of *Cyp7a1* and (G) *Cyp8b1* in WT and DKO females was lost in DKO females after OVX. (H) *Sult2a1* has greater baseline expression in DKO mice, induced to a lesser extent upon CA challenge compared to WT animals (*n* = 4-5/group). (I) ChIP-PCR was performed in WT and DKO male and female livers to test ERα recruitment to BA synthesis and metabolism genes, *Cyp7a1, Cyp8b1*, and *Sult2a1*. Mean ± SEM; Two-way ANOVA with Bonferroni post hoc analysis was performed. #p<0.05, *p<0.01, **p<0.001 compared to controls.

To overcome the confounding effects of ageing and cancer, we examined young WT and DKO female mice with and without OVX. Additionally, we challenged these mice with BA excess (Fig. 4D). As expected, OVX resulted in the reduction of basal hepatic *Erα* gene expression in both WT and DKO mice (Fig. 4E). In the DKO mice, which display high basal levels of BA synthesis and sulphation genes, we found dramatic induction of *Erα* gene upon BA treatment (Fig. 4E). Importantly, the rise in *Erα* gene coincided with reduced expression of *Cyp7a1, Cyp8b1* and increased levels of *Sult2a1,* a sulphotransferase known to sulphate estrogen and BAs. OVX in WT mice led to lower basal levels of *Cyp7a1 and Sult2a1 but not Cyp8b1,* whereas all three genes were significantly reduced in the DKO livers (Fig. 4G-4H). Unlike the OVX WT, which maintained CA-mediated suppression of BA synthetic genes, consistent with intact FXR signaling, DKO+OVX mice did not alter their expression (Fig. 4F-G). These data indicate a role for estrogen signaling in regulating BA homeostasis in the DKO livers.

We next examined if the recruitment of ERα to BA synthesis genes exhibited any sex difference in WT and DKO livers by ChIP-PCR. We find that ERα was preferentially recruited to *Cyp7a1* in a sex-specific manner (Figure 4I). *Cyp8b1* showed a similar trend but not *Sult2a1*. Also, we did not find any sex-specific patterns in ERα occupancy in *Ldlr* and *Pgr* genes, which were used as positive controls for ERα ChIP assays (Figure 4I). These data, along with increased BAs upon OVX, suggest ER*α* signaling is pertinent to control BA synthesis, especially in the absence of FXR, as seen in the CA-fed sham DKO mice.

### DKO mice display sexual dimorphism in BA homeostasis

WT mice do not show overt changes in serum BAs between the two sexes; however, genetically identical DKO mice displayed dramatically lower serum BAs in females compared to males (Supplementary Table 5). Nonetheless, the serum BA concentration in DKO females was higher than in WT females. This was intriguing. So, we analyzed the expression of genes involved in BA synthesis, transport, and metabolism in both sexes of DKO mice. Consistent with *Fxr* and *Shp* deletion that results in the loss of negative feedback on BA biosynthesis, both sexes of DKO mice have significantly higher expression of *Cyp7a1* and *Cyp8b1* genes that are involved in classical BA synthesis (Fig. 5A). The male dominant expression of *Cyp7b1* in the WT is lost in the DKO mice. On the other hand, *Cyp27a1*, which initiates alternative BA synthesis, was increased in a female-specific manner (Fig. 5A).

**Figure 5:**
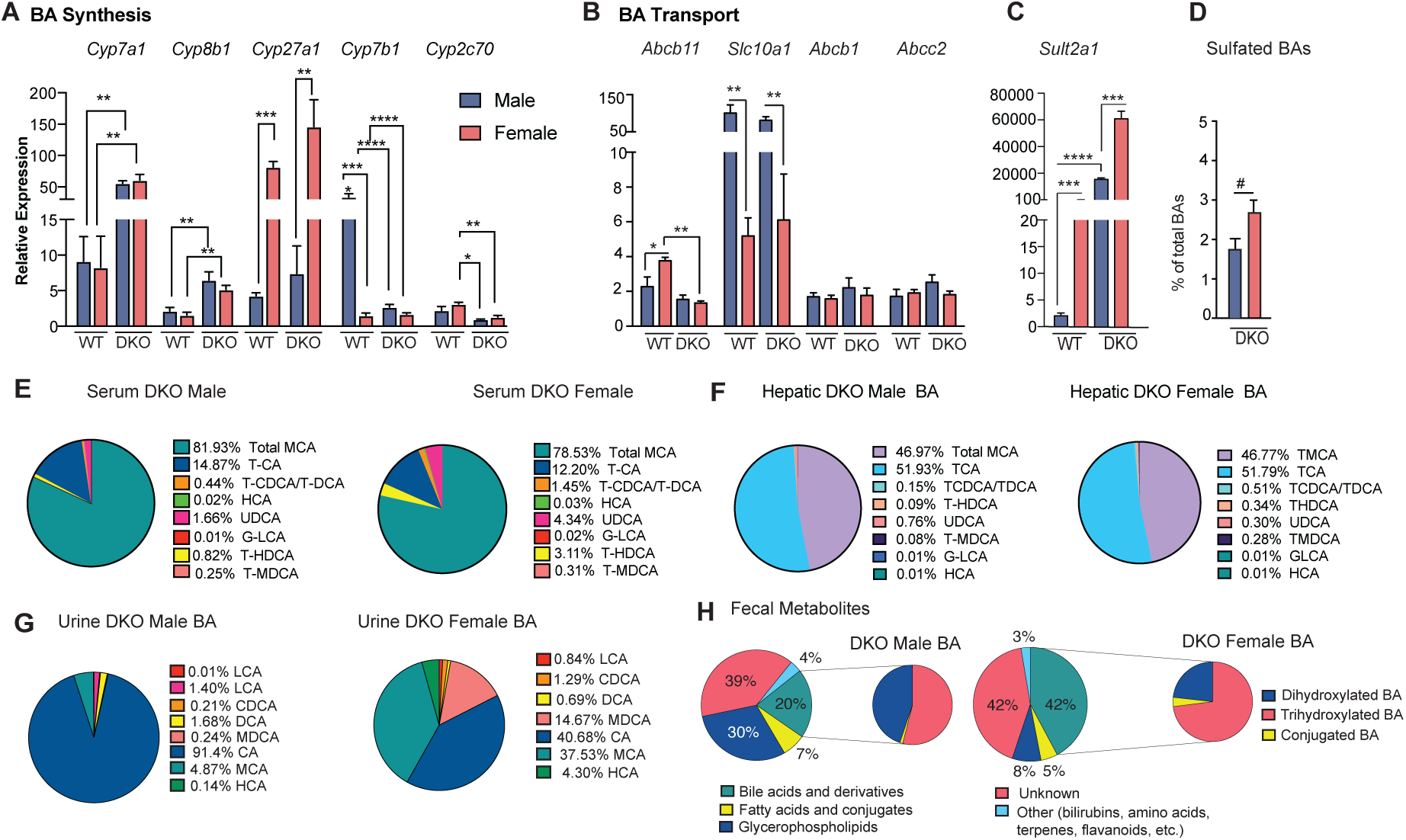
BA composition and metabolism are differentially regulated between the sexes of DKO mice. (A) Hepatic mRNA expression of classical BA synthetic enzymes was elevated in DKO compared to WT mice. While the alternative BA synthesis encoding gene, *Cyp27a1*, was increased in females only. (B) Expression of hepatic BA transporters and (C) BA sulfotransferase in WT and DKO mice. (D) Percentages of sulfated BAs in DKO male and female serum (one-tailed t-test). (E-F) BA composition is slightly varied in serum, whereas it remains unchanged in the liver between DKO males and females. (G-H) BA composition in the urine was variable between the sexes, and BAs constitute a higher proportion of fecal metabolites in the DKO females compared to males (*n* = 5-7/group). Mean ± SEM; #p<0.05, *p<0.01, **p<0.001 compared to genotype or gender controls. One-way ANOVA with Bonferroni post hoc analysis was performed.

Next, we examined BA transport. We found that hepatic transcript levels of the key BA efflux pump, bile salt export pump (*Bsep*, *Abcb11*), were reduced in both sexes of DKO mice, consistent with loss of *Fxr* (*24*) (Fig. 5B). In contrast, the expression of canalicular efflux transporters, *Abcb1 (Mdr1)* and *Abcc2 (Mrp2)* was unchanged (Fig. 5B). Also, the BA uptake transporter, sodium taurocholate co-transporting polypeptide (*Ntcp*, *Slc10a1*) showed lower transcript levels in females (Fig. 5B), which is in line with previous findings that estradiol can downregulate *Slc10a1* expression [54].

We then investigated the transcript expression of *Sult2a1*, which contributes to BA sulfation—a modification that can reduce enterohepatic recirculation [55]. As expected, hepatic *Sult2a1* expression was predominant in females irrespective of the genotype (Fig. 5C) [56]. Sulphated BAs are excreted in urine to eliminate excess BA during cholestasis [57, 58]. Total urine BA levels were higher in DKO males, reflecting a larger circulating BA pool than in DKO females (Supplementary Table 6). However, DKO female mice exhibited higher percentages of sulphated BAs (Fig. 5D), which corroborates with high *Sult2a1* expression in females.

BA compositional analysis was performed in the serum, hepatic, urine, and feces of DKO males and females (Fig. 5E-H). Both sexes of DKO mice showed abundant muricholates in the serum (Fig. 5E), but there were modest differences in the composition, indicating slightly hydrophilic BAs in the DKO females. Moreover, hepatic BA composition was indifferent between the two sexes (Fig. 5F). These results indicated that rather than synthesis or transport, excretion may be different between DKO males and females. Notably, we found that both urine and fecal levels and composition between male and female DKO mice were distinct (Fig. 5G-H). As urinary BA excretion alone cannot explain the 50% decrease in circulating BAs in DKO females, we performed untargeted metabolomics using the fecal samples. BAs accounted for 20% of the fecal samples in the males, whereas in the females, it was double the amount indicative of twice the amount being excreted in DKO females. These results indicate that female DKOs may be protected against detrimental tumor-promoting BA signaling due to their higher BA excretion.

### Increasing fecal BA excretion is sufficient to reduce liver cancer risk

Finally, to test this, we promoted fecal BA excretion in DKO males by using cholestyramine (CHR), a resin that binds BAs. We fed nine-month-old DKO male mice with a 2% CHR-containing diet since, by this age, tumor nodules have already developed. The CHR diet was continued until one year of age, mimicking a therapeutic intervention strategy (Fig. 6A). As expected, the CHR diet in DKO males led to a 50% reduction in circulating BA levels and altered BA composition (Fig. 6B-C). DKO males fed chow exhibited severe hepatic tumorigenesis, whereas CHR-fed DKO males had a drastically lower tumor burden with only small liver nodules and were protected from developing aggressive carcinomas. Histological analysis revealed that CHR treatment lowered the number of nodules and dysplastic changes but increased steatosis in DKO males (Fig. 6D). Conversely, increasing circulating BAs by causing biliary injury with 3,5-diethoxycarbonyl −1,4-dihydrocollidine (DDC) in DKO females resulted in the development of large liver tumors in DKO females (Supplementary Fig. 7).

**Figure 6:**
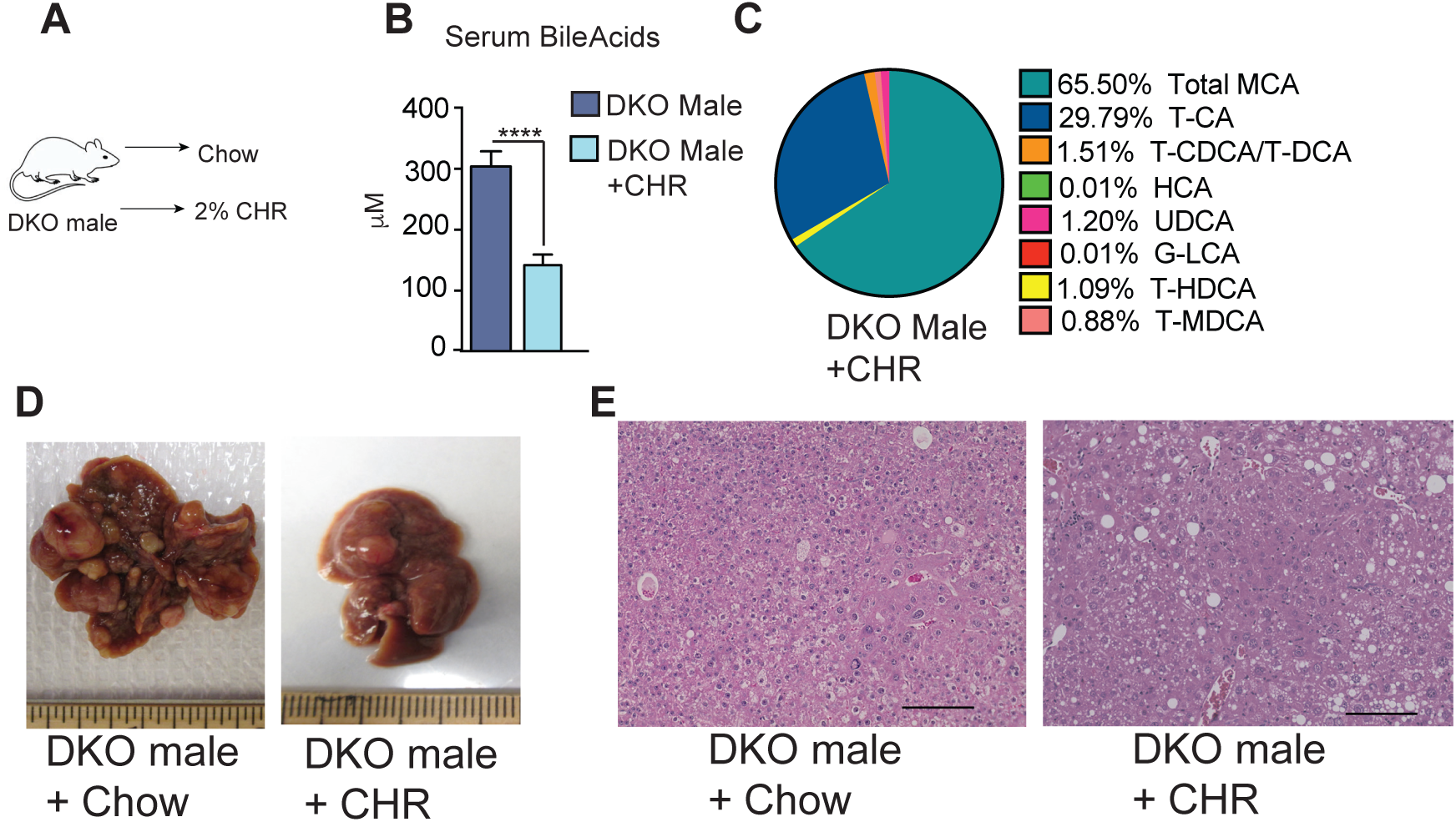
Treating with BA-binding resin reduces the tumor burden in DKO male mice. (A) DKO male mice were fed a 2% cholestyramine (CHR)-enriched diet for 3 months until one year of age. (B-C) Serum BA levels and composition upon feeding DKO male mice CHR-enriched diet. (D) CHR dramatically reduced the HCC burden in DKO males. (E) Histological analysis shows HCC, bland tumor cells, and enlarged nuclei with irregular membranes in DKO male mice. CHR treatment results in smaller and fewer nodules but increases steatosis. (*n* = 6-7/group). Mean ± SEM; *p<0.01; **p<0.001 compared to DKO controls.

Unbiased correlation analysis of hepatic and serum BA composition between the two cohorts of DKO mice revealed that the BA profiles of CHR-fed DKO male mice clustered with DKO females, whereas DDC-fed DKO female mice clustered with DKO males (Supplementary Fig. 8A).

Overall, these findings demonstrate that modulating circulating BAs is sufficient to change the liver cancer outcome, wherein lowering their levels leads to subsequent protection and vice versa.

## Discussion

Here, we demonstrate that the sex differences in BA homeostasis can contributes to the sexual disparity noted in HCC risk. Importantly, elevated BA concentrations are reported in patients with HCC [13, 26, 33–35, 59]. Using a genetic mouse model of excess BAs that develop spontaneous HCC, we uncovered distinct transcriptional control of metabolism between the two sexes. Both *Fxr* and *Shp* transcript levels were downregulated in HCC patients. Moreover, differential gene expression, specifically of the DKO female, correlated well with better survival, highlighting the translational relevance of our model. Thus, these gene signatures could be utilized as a potential prognostic marker for HCC progression and survival.

Both BA homeostasis and amino acid metabolism were altered between the two sexes. Of note, genes controlling ureagenesis were higher in the DKO females, and consistent with previous findings, we were able to recapitulate estrogen-mediated regulation of some of these genes signaling [48]. In line with these findings, patients with urea cycle enzyme deficiencies have a 200x higher incidence of HCC, highlighting the importance of amino acid metabolism in hepatic tumorigenesis [42–45]. Also, BAs have been shown to promote amino acid catabolic machinery [60], which indicates that BAs may be a central node in liver cancer. Intriguingly, hepatic urea analysis did not reveal any difference between the DKO male and female mice. A caveat being we measured a steady-state urea levels rather than the flux of this pathway.

We examined and found estrogen signaling can regulate the expression of BA synthesis and sulphation genes. DKO female mice challenged with the CA diet showed a robust increase in hepatic Erα transcript, which coincided with the suppression in BA synthesis in the absence of *Fxr* and *Shp*. Consistently higher recruitment of ER to the classical BA synthetic genes was noted in female livers. DKO OVX mice with blunted Erα gene expression, also showed lower transcript levels of *Cyp7a1 and Cyp8b1* that were unaltered post*-*CA challenge. Currently, we do not know how to reconcile this data other than indicating a potential ER-independent mechanism.

Nonetheless, we confirmed the known sex differences in BA synthesis, such as a female-dominant *Cyp27a1* expression and male-dominant *Cyp7b1* pattern in WT mice. Loss of *Fxr* and *Shp* altered the expression of many genes irrespective of sex. For instance, C*yp2c70* expression was reduced in *both sexes*, and the male dominance of *Cyp7b1* was lost in the DKO mice.

Of note, OVX of DKO females increased the serum BA levels and lost their protection against the development of liver tumorigenesis. This finding fully recapitulates the clinical data, wherein post-menopausal women are equally prone to HCC incidence as males.

BA analysis shows that DKO female mice have a hydrophilic composition and excrete BA proportions. So, we tested and demonstrated the potent therapeutic utility of reducing BA levels in serum using a generic FDA-approved BA binding resin, Cholestyramine (CHR), in dramatically reducing the tumor burden. This study highlights that lowering enterohepatic recirculation is a beneficial strategy in modulating liver cancer. Though *Cyp7a1* expression is reported to be induced in CHR-fed mice [61], long-term CHR feeding in DKO mice lowered *Cyp7a1* expression but induced *Cyp8b1* transcripts (Supplementary Figure 8B). Conversely, DDC-fed DKO females that develop hepatic tumors show a corresponding decrease in *Cyp8b1* transcript (Supplementary Figure 8C). Also, patients with HCC exhibit a reduction in *Cyp8b1* expression [62–64], which promotes a more hydrophilic ratio of BA composition.

Although species differences in BAs between mice and humans are a limitation, several fundamental understandings have been gained from mouse experiments. While this study demonstrates bile acids promote HCC progression, it does not investigate or provide evidence if excess bile acids are sufficient for HCC initiation. Another caveat is that the DKO mouse model mimics the progression of cholestasis to HCC and not all the etiologies, so the observed sex differences in circulating BAs may be limited to these subsets of HCC. Nevertheless, elevated BA concentrations are seen in various liver disease conditions and inherited disorders of cholestasis predispose to HCC onset. More recently, clinical studies support the utilization of BAs as prognostic markers [65–67].

Our findings demonstrate that female cholestatic mice exhibit increased excretion and lower serum BAs than males. However, hepatic BAs were not different between the sexes. These data highlight that circulating BAs contribute towards sex differences seen in HCC incidence. Accordingly, we show that lowering enterohepatic BA recirculation is beneficial in our model. Our results align with previous findings that had implicated intestinal FXR signaling as being crucial rather than hepatic FXR to prevent liver tumorigenesis [68]. Taken together, these results reveal that drugs inhibiting intestinal reabsorption of BAs (Asbt inhibitor, IBAT inhibitor) that are on clinical trials for NASH and cholestasis can be evaluated as potential therapeutics to combat HCC.

## Methods

### Experimental Design

This study was designed to elucidate the role of bile acids (BAs) in the sexually dimorphic incidence of HCC and assess the therapeutic benefits of reducing circulating BA levels on HCC development. Fxr^-/-^, Shp^-/-^ (DKO) mice were maintained on a C57BL/6 background at the University of Illinois, Urbana-Champaign. Male and female mice were sacrificed at 8-12 weeks or 6 and 12 months after birth. Mice were housed on a standard 12-hour light/dark cycle and fed normal chow and water ad libitum. In order to study estrogen signaling, bilateral ovariectomies were performed on WT and DKO females at 8-10 weeks old, followed by one week of recovery. 2% Cholestyramine-supplemented chow was fed to 9-month-old DKO male mice for a period of 3 months while 0.1% DDC (3,5-diethoxycarbonyl-1, 4-dihydrocollidine) was fed to 10-month-old DKO female mice for 3 months. Urea cycle studies were performed on mice after overnight fasting. All studies were carried out as outlined in the Guide for the Care and Use of Laboratory Animals prepared by the National Academy of Sciences and published by the National Institutes of Health (National Institutes of Health publication 86-23, revised 1985). For biological harvesting, mice were anaesthetized and euthanized as described by IUCAC. Tissue was flash-frozen in liquid nitrogen and blood was collected and spun down for serum.

### Serum Chemistry

Blood was collected by retro-orbital bleeding and centrifuged at 8000g x 10 minutes to separate serum. Serum ALT and AST were measured using Infinity ALT and Infinity AST kits (Thermo Scientific). Calorimetric measurement of serum and hepatic bile acids was performed with the Total Bile Acid (NBT method) kit (Genway Biotech).

### Bile Acid analysis

Serum and urine from DKO male and female mice fed chow were analyzed for the composition of bile acids and their sulfated metabolites at the University of Nebraska Medical Center. Liquid chromatographic-mass spectrometry analysis was performed with a Waters ACQUITY column (Milford, MA). Bile acids and internal standards were prepared in methanol and analyzed. Serum from DKO male and female mice fed chow, DKO males fed CHR, and DKO females fed DDC was analyzed for bile acid composition at Baylor College of Medicine Metabolomics Core, Houston, Texas. Liquid chromatographic-mass spectrometry analysis was performed with a Waters ACQUITY UPLC BEH C18 column (Milford, MA). Bile acids were detected in negative mode, with L-Zeatine added to each sample as an internal standard.

### Metabolite Profiling

Liver tissue was weighed and sonicated in 70% methanol, followed by centrifugation. The supernatant was flash-frozen and used for subsequent LC-MS analysis for urea cycle metabolites. Tissue lysate was used for the BCA assay to determine the protein concentration of each sample. All metabolite concentrations were normalized to a protein concentration of the lysate.

### Untargeted Metabolomics

Fecal samples were weighed into microcentrifuge tubes and homogenized in 50% MeOH/H2O solution with a 1:10 w/v ratio, for 5 minutes at 5 Hz. The samples were centrifuged at 14000 rpm for 15 minutes, then a 200uL aliquot of each supernatant was transferred to a 96-well plate and dried under centrifugal vacuum. The dried extracts were covered and stored at −80 °C until analysis, at which time the samples were resuspended in 200uL of 50% MeOH/H2O solution with 1uM sulfadimethoxine as internal standard and diluted three-fold for analysis. Untargeted LC-MS/MS was performed on a Thermo Vanquish UPLC system coupled to a Q-Exactive Orbitrap mass spectrometer (ThermoFisher Scientific, Bremen, Germany). A polar C18 column (Kinetex polar C18, 100 x 2.1 mm, 2.6 μm particle size, 100 A pore size; Phenomenex, Torrance, CA USA) was used as the stationary phase, and a high-pressure binary gradient pump was used to deliver the mobile phase, which consisted of solvent A (100% H2O + 0.1 % formic acid (FA)) and solvent B (100% acetonitrile (ACN) + 0.1 % FA). The flow rate was set to 0.5mL/min and the injection volume for each sample was 5uL. Following injection, samples were eluted with the following gradient: 0-1.0 min, 5% B; 1.0-1.1 min, 25%; 6.0 min, 70%; 7.0 min, 100%; 7.5-8.0 min, 5%. MS data was collected in positive mode and electrospray ionization (ESI) parameters were set to 53 L/min for sheath gas, 14 L/min for auxiliary gas, 0 L/min for spare gas, and 400°C for auxiliary gas temperature. The spray voltage was set to 3500 V, the capillary temperature to 320 °C, and the S-Lens radio frequency level to 50 V. MS1 data were collected from 150-1500 m/z with a resolution of 35,000 at m/z 200 with one micro scan. The maximum ion injection time was set to 100 ms with an automatic gain control (AGC) target of 1.0E6. MS/MS spectra were collected using data dependent acquisition (DDA), where the top 5 most abundant ions in the MS1 scan were selected for fragmentation. Normalized collision energies were increased stepwise from 20, 30, to 40. MS2 data were collected with a resolution of 17,500 at m/z 200 with one micro scan and an AGC of 5.0E5. All untargeted LC-MS/MS data are publicly available from the MassIVE data repository under accession number MSV000089715. MS1 feature detection and MS/MS pairing was performed using MZmine 2.37corr17.7_kai_merge (55, 56). An intensity threshold of 5E4 and 1E3 were set for MS1 and MS2 detection, respectively, with centroid data. MS1 chromatogram construction was performed using the ADAP chromatogram builder, where the minimum group size was set to 5, group intensity threshold was 5E4, minimum highest intensity was 1.5E5, and mass tolerance was 0.005 m/z or 10 ppm. Chromatogram deconvolution was then performed using a local minimum search algorithm with a chromatographic threshold of 80%, a search minimum in retention time (RT) range of 0.2 min, minimum relative height of 1%, minimum absolute threshold height of 1.5E5, minimum ratio for top/edge of 1, and a peak duration of 0.05-2.0 min. Pairing between MS1 and MS2 was performed with a mass tolerance of 0.005 m/z or 10 ppm and RT range of 0.2 min. Isotope peaks were grouped, then features from different samples were aligned using the same mass and RT tolerances; alignment was performed by placing a weight of 75 on m/z and 25 on RT. A peak area feature table was exported as a .csv file and consensus MS/MS spectral data were exported in mgf format. Feature-based molecular networking and MolNetEnhancer workflows were then performed with this data using GNPS (gnps.ucsd.edu). The corresponding jobs can be found at: https://gnps.ucsd.edu/ProteoSAFe/status.jsp?task=d697d44ec18440d29d0771f84ba7cccdand https://gnps.ucsd.edu/ProteoSAFe/status.jsp?task=e3edba56efba4a27b073a9031c60b5e5, respectively.

### Histology

Liver samples were fixed in 10% neutral-buffered formalin, sections were cut at a thickness of ∼5 microns and used for hematoxylin and eosin, and sirius red staining.

### RNA extraction and quantitative PCR analysis

Total RNA from the liver was prepared according to the TRIzol (Invitrogen) protocol. cDNA was synthesized using Maxima Reverse Transcriptase (Thermo Scientific) according to the manufacturer’s protocol. q-RTPCR was performed on an Illumina Eco Platform. For qRT-PCR analysis, 50 ng of cDNA was added to each SYBR green-based reaction. qRT-PCR primers are provided in Supplementary Table 7.

### Microarray

Microarray was performed by Dr. Ju-Seog Lee’s laboratory at the MD Anderson Cancer Center. Liver samples from 12-month-old male and female WT and DKO mice were collected and snap-frozen. Total RNA was isolated, labeled, and hybridized to BeadChip Array MouseWG-6 (Illumina). Bead chips were scanned with an Illumina BeadArray Reader. Microarray analysis was performed on the Illumina mouseRefseq-8 Expression platform. Upregulated gene sets were generated from genes with fold change > 1.3 (p<0.0001) compared to the control group (i.e. DKO males vs. DKO females). These gene sets were then used for downstream analyses with DAVID Bioinformatics Resources Analysis Software and ClueGO [36].

### Transcription Factor Motif Analysis

GeneXplain software was used to identify enriched transcription factor binding sites (TFBS) using the upregulated gene sets generated from the microarray. The analysis included regions from −1000 to 100 bp relative to the transcription start site. Only those TFBS enriched with p≤ 0.01 were included in the tables.

### Extraction of transcriptomic signature

Multiple transcriptomic signatures were extracted from the microarray data of the DKO mouse model (Supplementary Table 1). DKO_All signature was generated from the comparison between wild type (WT) male and female mice, and DKO_Male and DKO_Female signatures from WT male and female mice, respectively. DKO_FvsM, DKO_Estrogen, DKO_BA, and DKO_Urea signatures were made from the comparison between DKO male and female mice. Signature genes were selected by T-test and logFC (p<0.001 and log2FC>1 or <-1) using the gene expression dataset after normalization.

### Gene expression data from HCC tumors

Gene expression data from the National Cancer Institute (NCI) cohort were generated in earlier studies [69–71], and the data are publicly available from the NCBI’s GEO database (GSE1898 and GSE4024). Gene expression data from Korea, Samsung, Modena, and Fudan cohorts have been described previously and are available from the NCBI’s GEO database (accession numbers, GSE14520, GSE16757, GSE43619, GSE36376, and GSE54236) [72–76]. TCGA RNA sequencing data for HCC was downloaded from the University of California, Santa Cruz, Genomics Institute (https://xenabrowser.net/) 12. FPKM-normalized data were log-transformed.

Tumor specimens and clinical data were obtained from HCC patients who had undergone hepatectomy as a primary treatment for HCC at multiple institutes, as described in their original study. Except for the TCGA cohort, patients and tissues were collected based on the availability of high quality of frozen tissues for genomic studies. For TCGA cohort [77], surgical resection of biopsy biospecimens were collected from patients diagnosed with HCC and had not received prior treatment for their disease (ablation, chemotherapy, or radiotherapy). Institutional review boards at each tissue source site reviewed protocols and consent documentation and approved the submission of cases to TCGA. Hematoxylin and eosin (H&E) stained sample were subjected to independent pathology review to confirm that the tumor specimen was histologically consistent with the allowable HCC. Each case was reviewed independently by at least 3 liver pathologists, with no clinical or molecular information.

### Prediction model

To predict the class similar to the DKO signature in the human HCC cohort, we used a classification algorithm based on Bayesian compound covariate predictor (BCCP). After the integration of the signature matrix and the human HCC dataset, the Bayesian probability for each human HCC sample were calculated by using the class prediction procedure in BRB Arraytools. We set 0.5 as the cut-off of Bayesian probability for each signature.

### Validation of clinical relevance

Prognostic significance was evaluated rigorously for overall and recurrence-free survival in the human HCC cohort based on the predicted class calculated by the BCCP algorithm using multiple DKOsignatures. A total of 5 human HCC transcriptomic cohorts were used in this study (Fudan, Korea, Samsung, TCGA, Modena). All DKOsignatures were evaluated in each human HCC cohort and meta-cohort. To identify the gender difference in the human HCC cohort, we did subgroup analysis for gender and age in the meta-cohort. BCCP scores (BCCP probability) were compared in all populations and gender subgroups. The analysis for potential correlation between the class predicted by DKO-signature and staging HCC in terms of TNM, BCLC, and CLIP classification was performed in the meta-cohort.

### ERα ChIP-Seq Analysis

ERα-ChIP assay was performed in both sexes of WT and DKO mice. ERα-F10 antibody (sc-8002, Santa Cruz) was used to perform the pulldown followed by qPCR. We also analyzed BED files for ERα ChIP-Seq from three independent studies [78–80], which were obtained from Cistrome DB and visualized using the UCSC genome browser on mouse GRCm38/mm10 assembly. For tracks 3 and 4, the track size was set to auto-size.

### Statistical Analysis

All statistical tests were performed using GraphPad Prism software. Data are presented as means ± SEM. Multiple group comparisons were analyzed using one-way and two-way ANOVA with the post hoc Bonferroni test. Unpaired t-test was used for comparison between two groups. P values ≤0.05 were determined to be significant unless otherwise noted in legends.

## Data Availability

The gene expression data generated and used in this publication have been deposited in NCBI’s Gene Expression Omnibus and are accessible through GEO Series accession number GSE151524. (https://www.ncbi.nlm.nih.gov/geo/query/acc.cgi?acc=GSE151524).

## Acknowledgments

We thank the Systems Biology laboratory at the University of Texas, MD Anderson Cancer Center, for the initial analysis and for performing the microarray studies. We would also like to thank the metabolite analysis core at Baylor College of Medicine for performing the bile acid composition analysis. We thank Dr. Bhoomika Mathur for her initial help with the DDC experiment. The authors also thank Drs. Auinash Kalsotra and Stephanie Ceman for their comments and critiques during the preparation of this manuscript. We also thank Ms. Angela Major at Texas Children’s’ Hospital for histological preparation and analysis. The authors thank Dr. Lucas Li at the Roy Carver Metabolomics Core at the University of Illinois at Urbana-Champaign for Urea Cycle metabolite analysis.

## Author contributions

M.E.P., S.K., A.E.D and S.A. contributed to conception, experimental design, data acquisition, analysis and interpretation, and drafting the article. L.J.T. helped with urea cycle expression analysis, R.N.T. and Y.A. analyzed bile acid composition and interpretation from serum and urine, E.G., M.P, P.D performed untargeted metabolomics, A.E.D, Q.Z, Z.M.E analyzed ERa ChIP-seq data, H.S.L and J.S.L performed all the HCC patient cohort analysis, M.J.F. evaluated and interpreted the liver histology.

---

**Supplementary Fig. 1:**
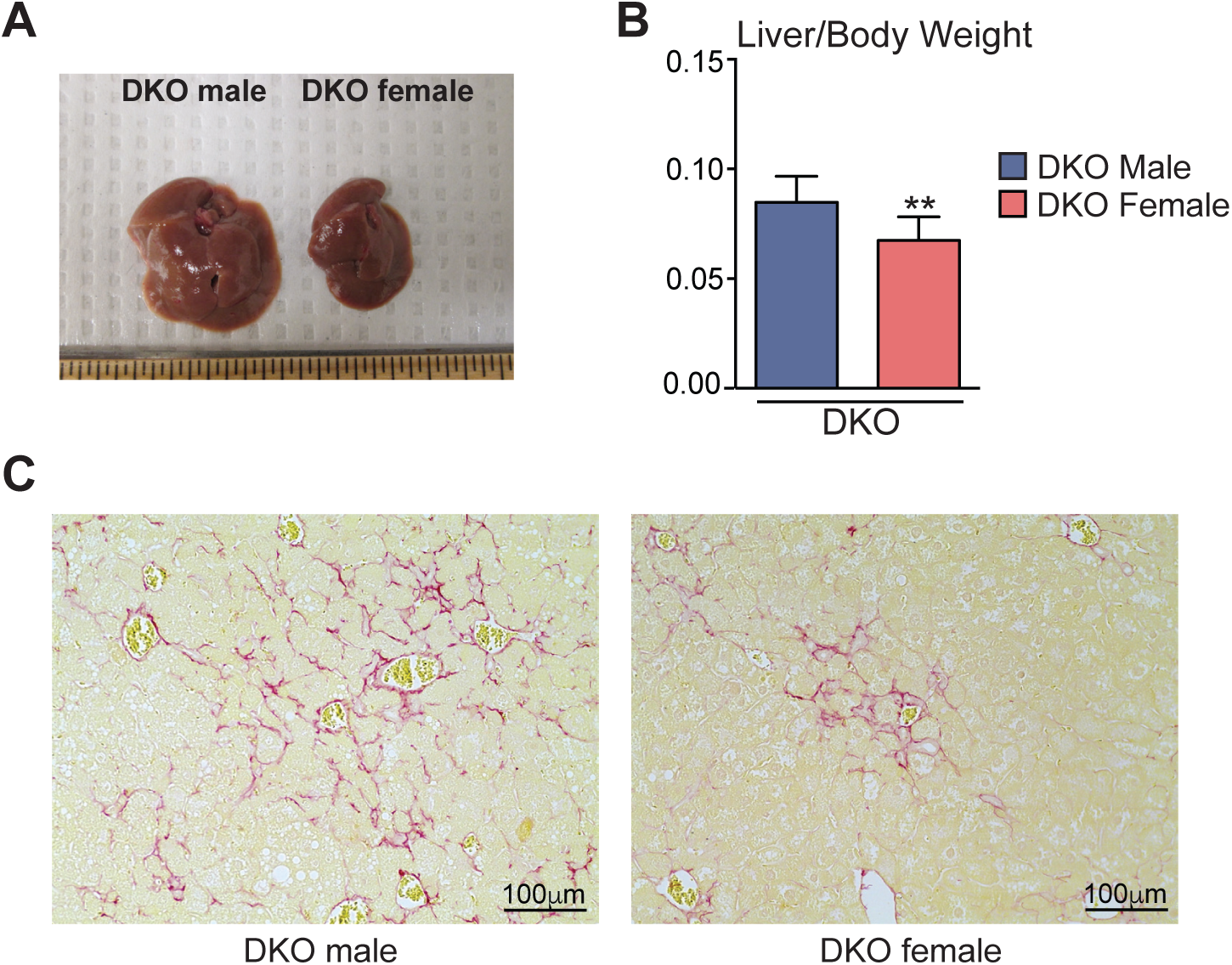
DKO female mice exhibit reduced liver injury. (A-B) Compared to DKO males, the female mice showed smaller liver size and liver-to-body weight ratio. (C) Sirius red staining revealed less collagen staining (red) in DKO females. (Six month old mice, n = 6/group). Mean ± SEM. *p<0.01, **p<0.001 compared to genotype or gender controls. Unpaired t-test was used to analyze the data.

**Supplementary Fig. 2.**
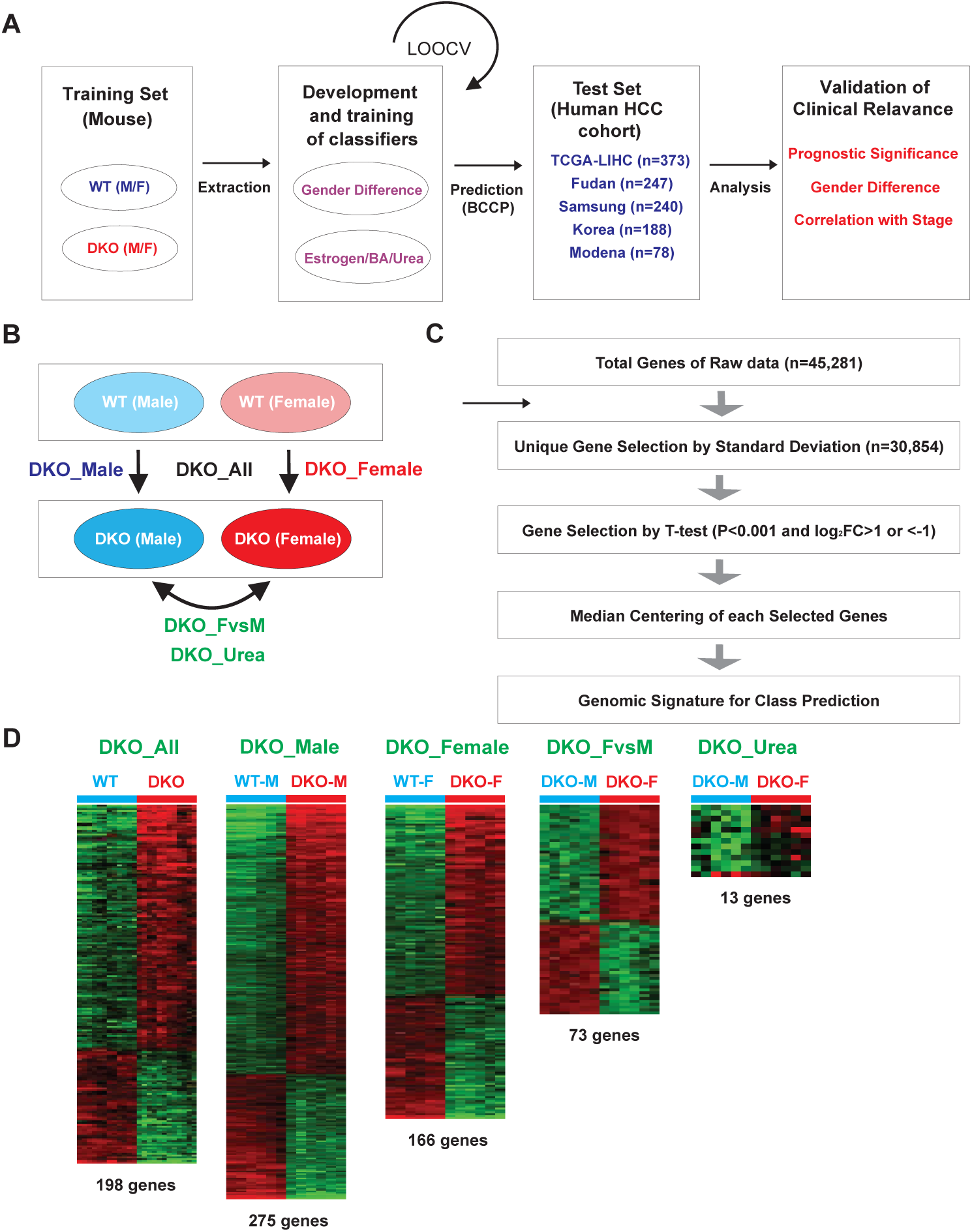
Schematic of the pipeline used to obtain gene signatures to predict outcomes in HCC patient cohorts. (A) Workflow schematic of the gene expression analysis. (B-C) Different gene signatures were extracted from the microarray data by comparing wild-type (WT) male and female gene expression patterns with that of DKO male and DKO female, respectively. These gene lists were used to predict overall survival and recurrence-free survival using five human clinical cohorts. (D) Gene changes in these defined sets DKO males (DKO_M), DKO females (DKO_F), or combined (DKO_ALL), or DKO_F vs M (DKO female gene signature that does not overlap with DKO males), DKO_Estrogen, DKO_BA, and DKO_Urea are shown. These genes were selected by t-test and log2 <u>F</u>old <u>C</u>hange (p<0.001 and log2 FC>1 or <-1).

**Supplementary Fig. 3:**
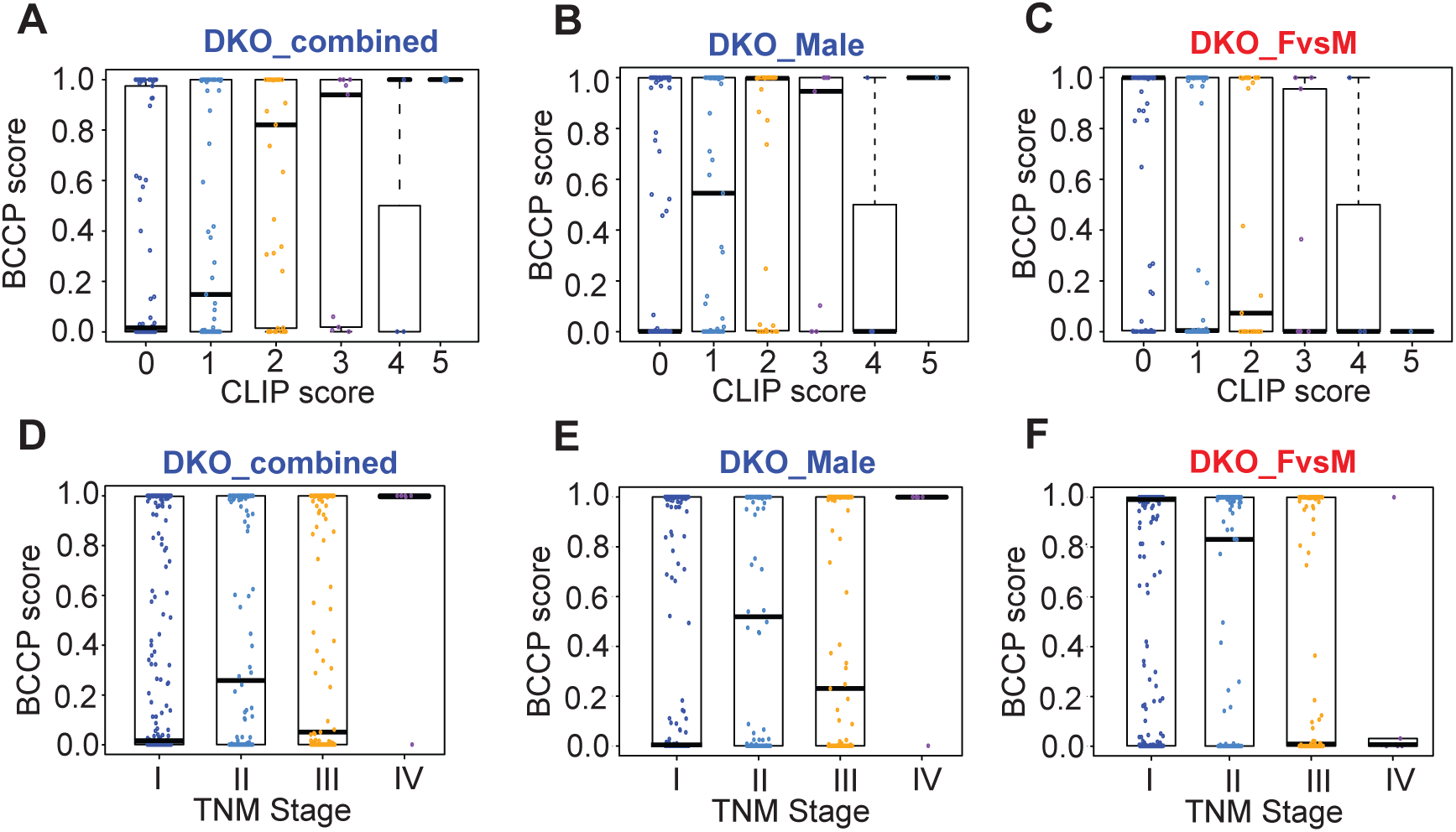
Correlation of DKO gene signatures with various clinical stages of HCC. Clinical relevance of combined gene changes in DKO male and female (DKO_Combined), or DKO male-specific, and the distinct gene set of DKO female vs. male (DKO_FvsM) were tested in five HCC cohorts. The trained BCCP (Bayesian compound covariate predictor) algorithm using the three DKO gene sets was tested across the HCC stages as determined by (A-C) CLIP (Cancer of the Liver Italian Program) or (D-F) TNM (Tumor, Node, and Metastasis). The probability of gene signatures was generated from 0 (not predictable) to 1 (predictable).

**Supplementary Fig. 4:**
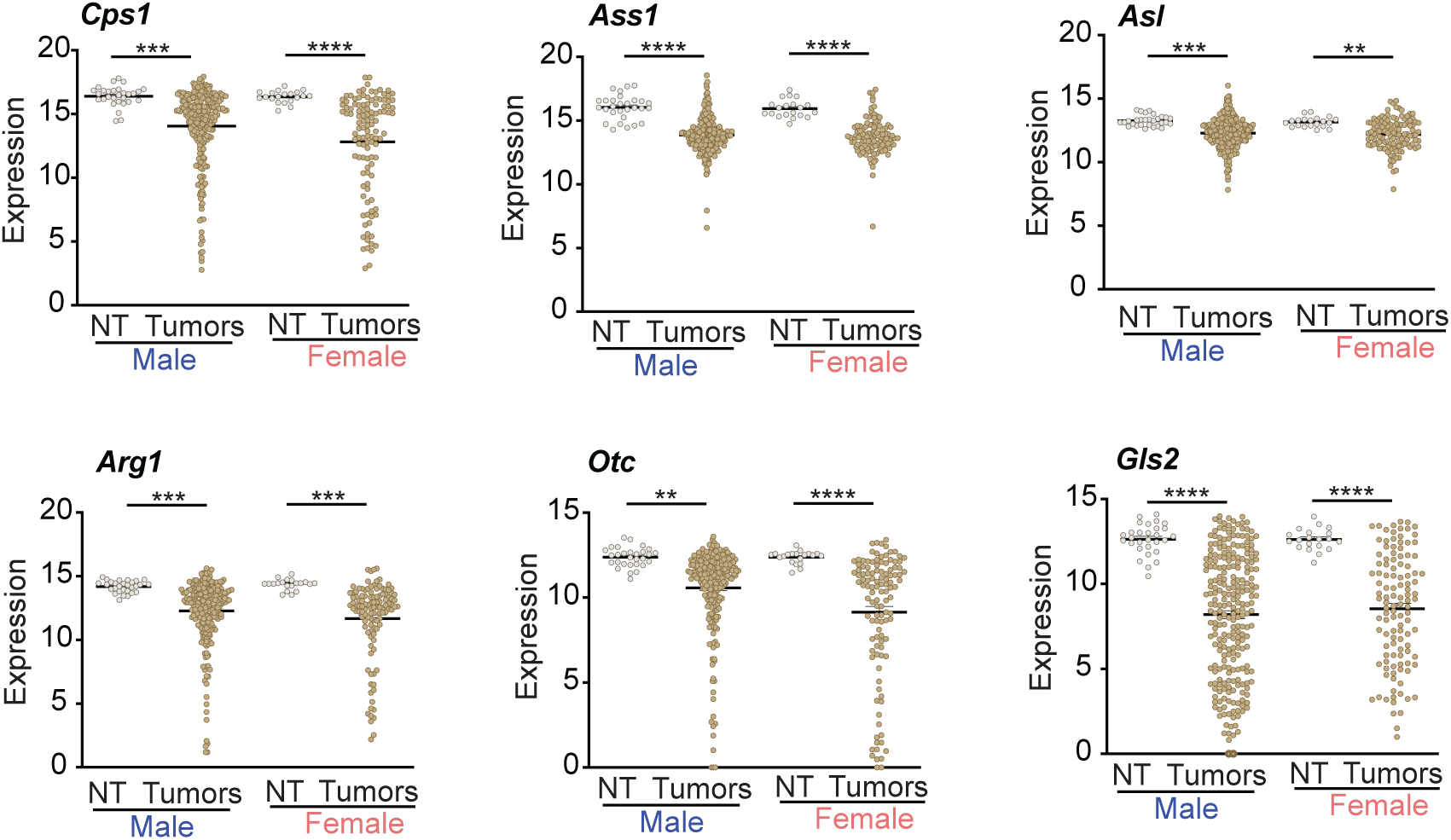
Expression of the urea cycle genes are reduced in human liver cancer. RNA-seq from TCGA-LIHC (The Cancer Genome Atlas-Liver Hepatocellular Carcinoma Collection) database was analyzed. Several genes that encode enzymes involved in ureagenesis exhibited a reduction in their transcript levels in tumors compared to non-tumor (NT) tissue. (Males: n =29 NT, n=245 tumors; Females: n=20 NT and n=114 tumors). Mean ± SEM; *p<0.01, **p<0.001, ***p<0.0001 compared to their respective sex-specific controls. One-way ANOVA with Bonferroni post hoc analysis was performed.

**Supplementary Fig. 5:**
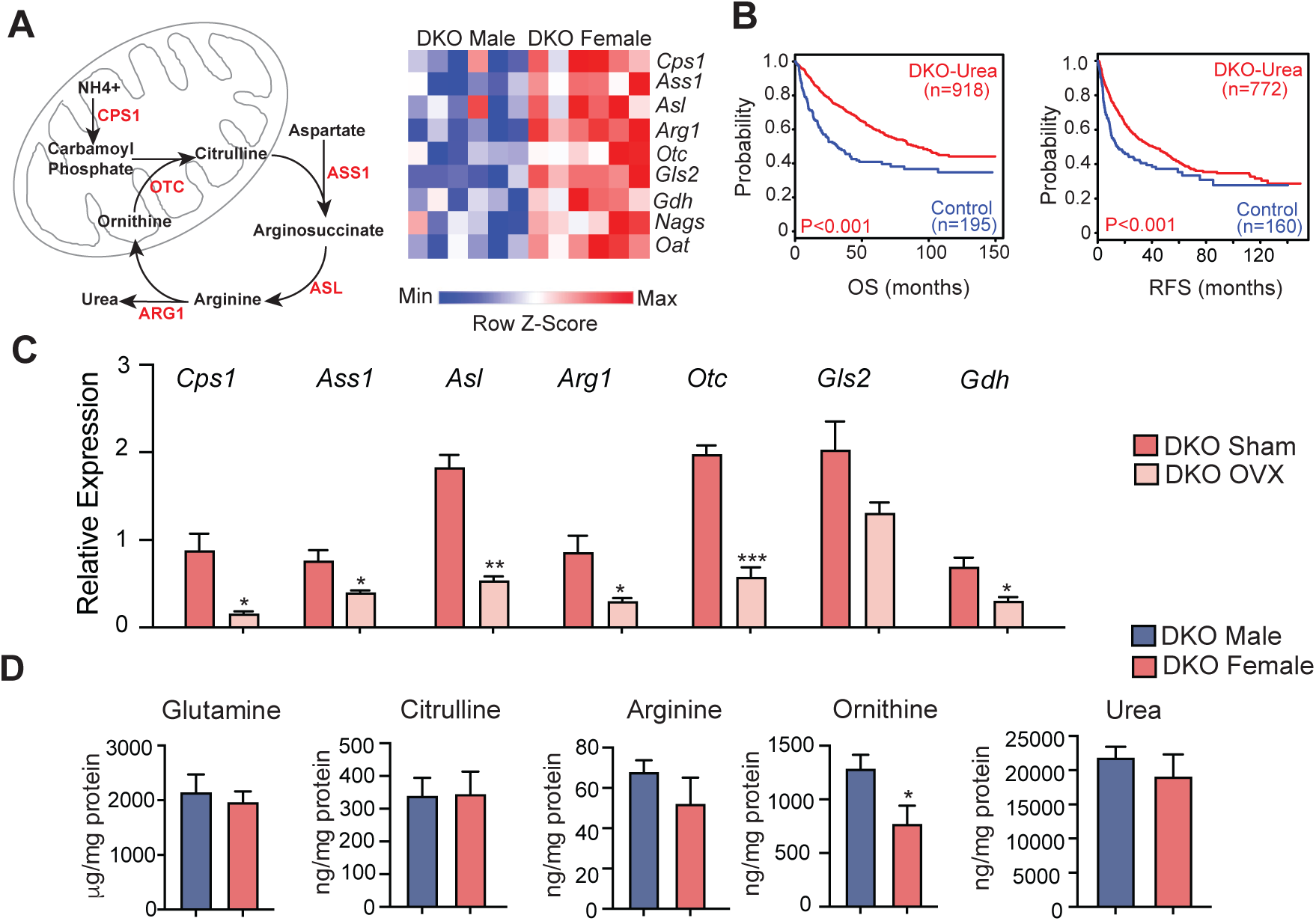
Urea cycle genes elevated in DKO females correlate with better patient survival and reduced after ovariectomy. (A) Schematic of the urea cycle, with a representative heat map of urea cycle genes from the microarray showing higher expression in DKO female livers (n = 6/group). (B) Survival curves from HCC patients who exhibit increased expression of urea cycle genes. (C) Expression of urea cycle genes was reduced in DKO female mice following OVX (*n* = 4-5/group). and (D) LC-MS quantification of urea cycle intermediates (*n* = 4-5/group). Mean ± SEM; *p<0.01, **p<0.001 compared to genotype or gender controls. Unpaired t-test was used for analysis.

**Supplementary Fig. 6:**
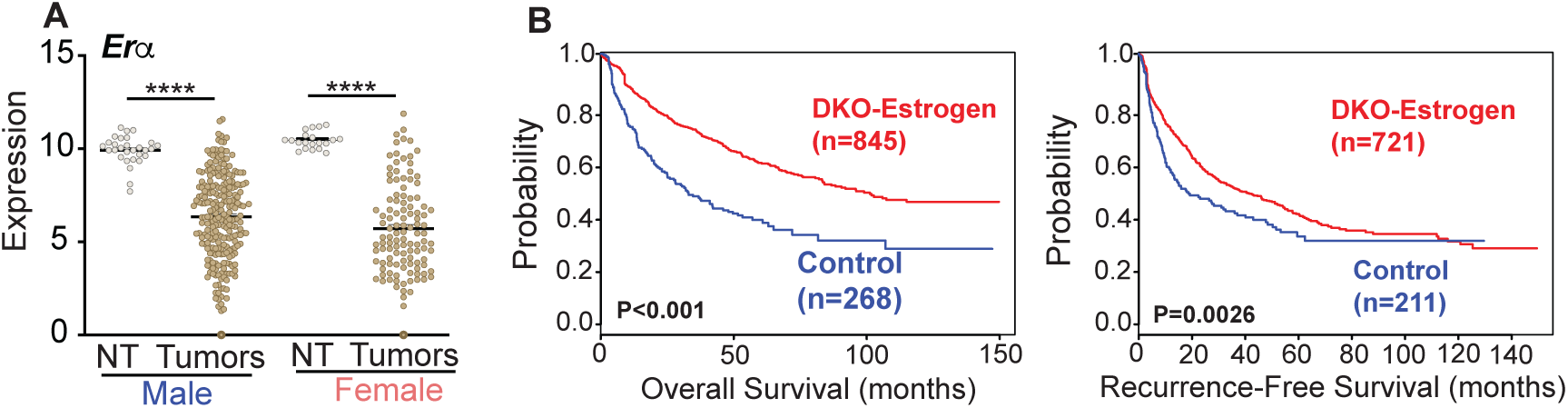
Estrogen receptor signaling positively correlates with better survival in HCC clinical samples. (A) RNA-seq from TCGA-LIHC (The Cancer Genome Atlas-Liver Hepatocellular Carcinoma Collection) data was analyzed. *Er*α transcript levels were reduced in tumors compared to non-tumor (NT) tissue. (Males: n =29 NT, n=245 tumors; Females: n=20 NT and n=114 tumors). Mean ± SEM; *p<0.01, **p<0.001, ***p<0.0001 compared to their respective sex-specific controls. One-way ANOVA with Bonferroni post hoc analysis was performed. (B) Kaplan Meier Survival graphs using Erα targets altered in DKO females (DKO_Estrogen) showed better clinical outcomes for HCC patients.

**Supplementary Fig. 7:**
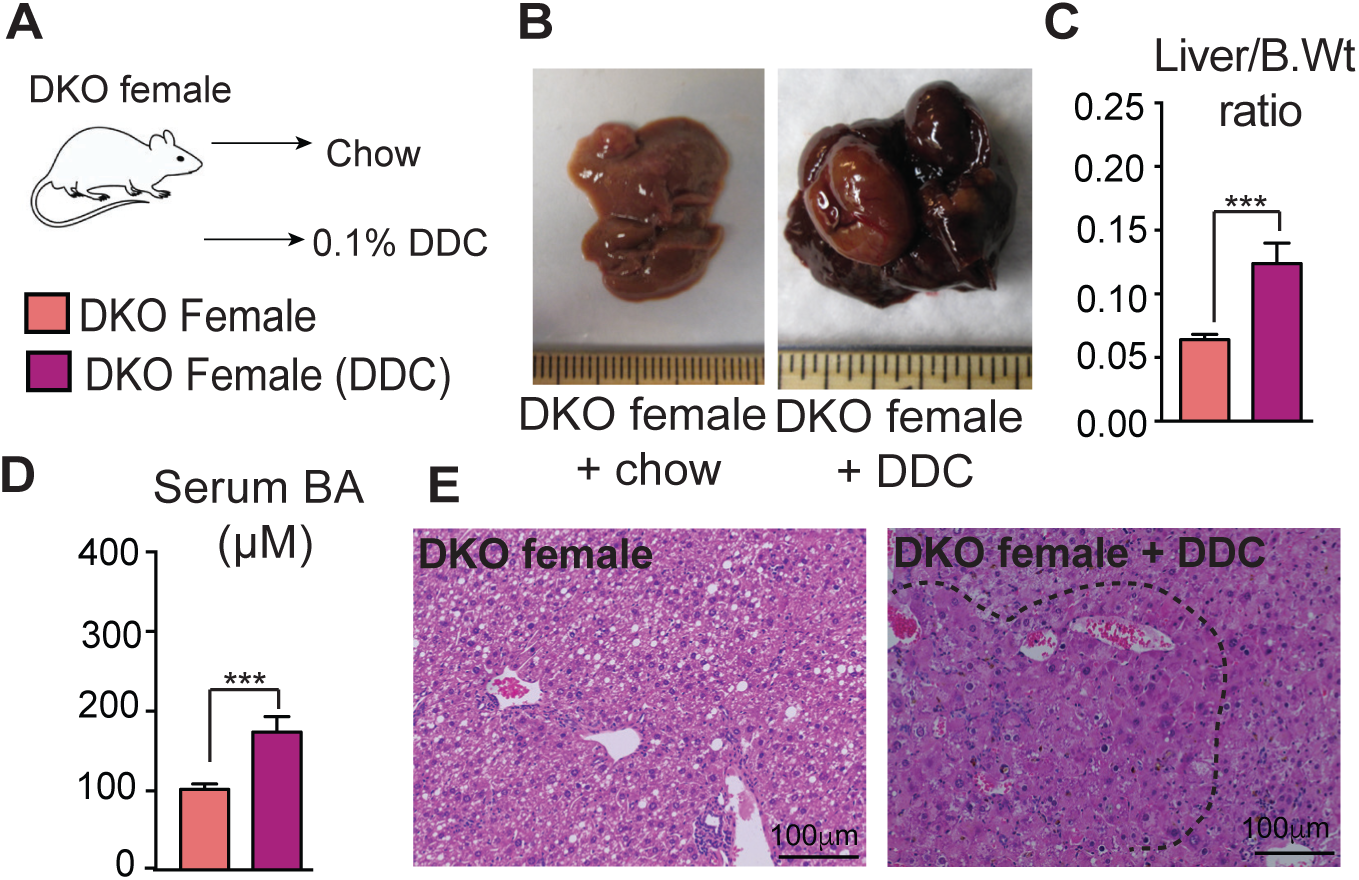
DKO females challenged with the DDC diet developed liver tumors. (A-B) DKO females were fed a 0.1% DDC diet, as shown in the schematic, for 5 months, and this treatment led to visible tumors at one year. (C-D) The diet elevated liver-to-body weight ratio and serum BAs in DKO females. (E) Histology revealed cholangitis, ductular reaction, and tumor nodules (marked by a dotted line) in DKO female livers+ DDC treatment. (*n* = 4-7/group). Mean ± SEM; *p<0.01; **p<0.001 compared to chow-fed DKO controls. Unpaired t-test was used for analysis.

**Supplementary Fig. 8:**
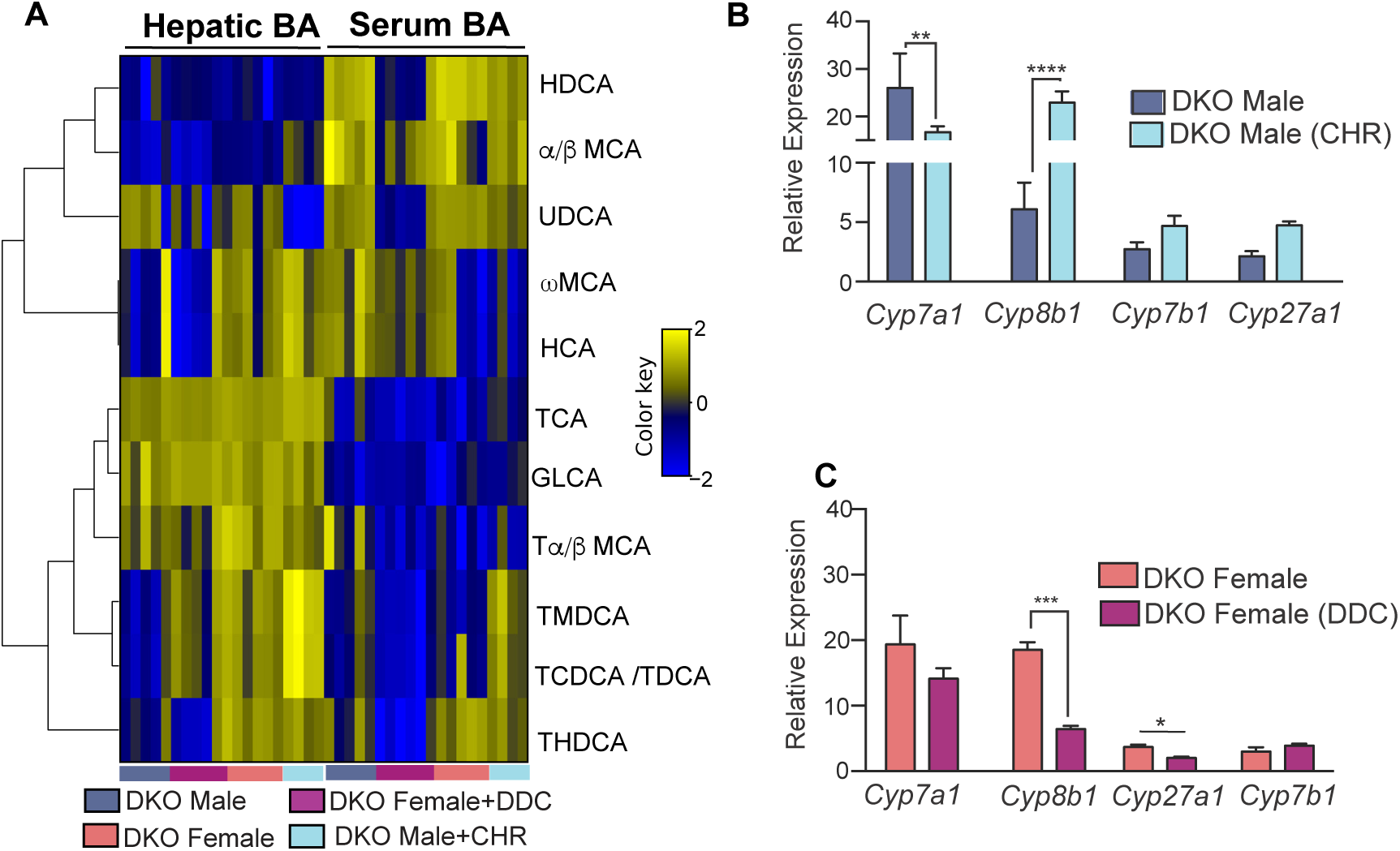
BA composition and expression of BA synthesis genes in DKO mice after different treatments. (A) Correlation analysis of serum and hepatic composition of BAs post-DDC diet in DKO female mice (n = 5-8 mice/group) and post-CHR diet in DKO male mice (n = 4-5 mice/group) along with their sex-specific chow controls. (B-C) Relative expression of classical and alternative BA synthesis genes after CHR and DDC treatments. Mean ± SEM; *p<0.01; **p<0.001, *** p<0.0001 compared to their chow-fed DKO controls. Unpaired t-test was used for analysis.

**Supplementary Table 1:**

**List of genes in different DKO gene signature category used for analysis See Excel sheet**

**Supplementary Table 2.**
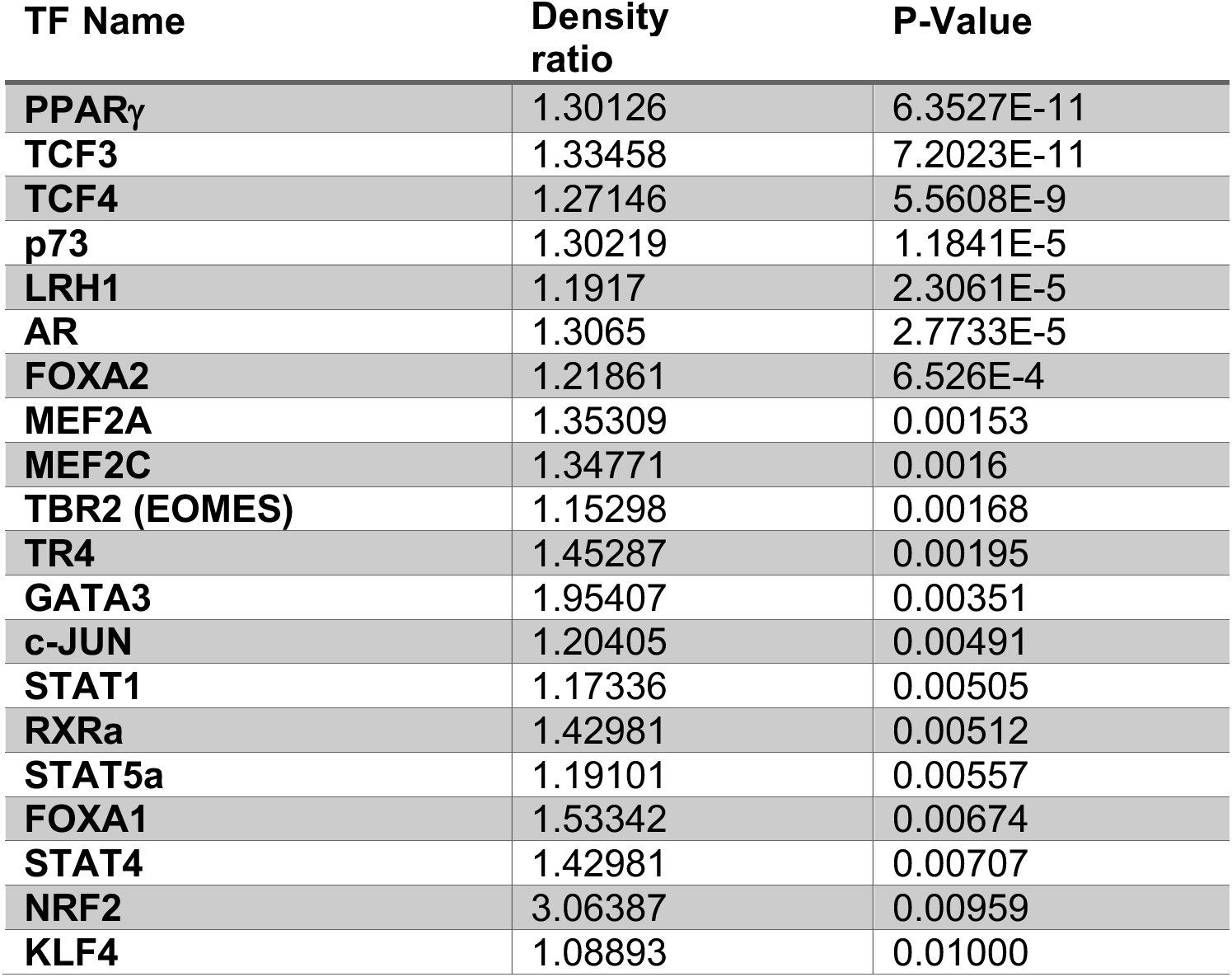
Transcription Motifs Enriched in DKO male livers compared to DKO females.

**Supplementary Table 3.**
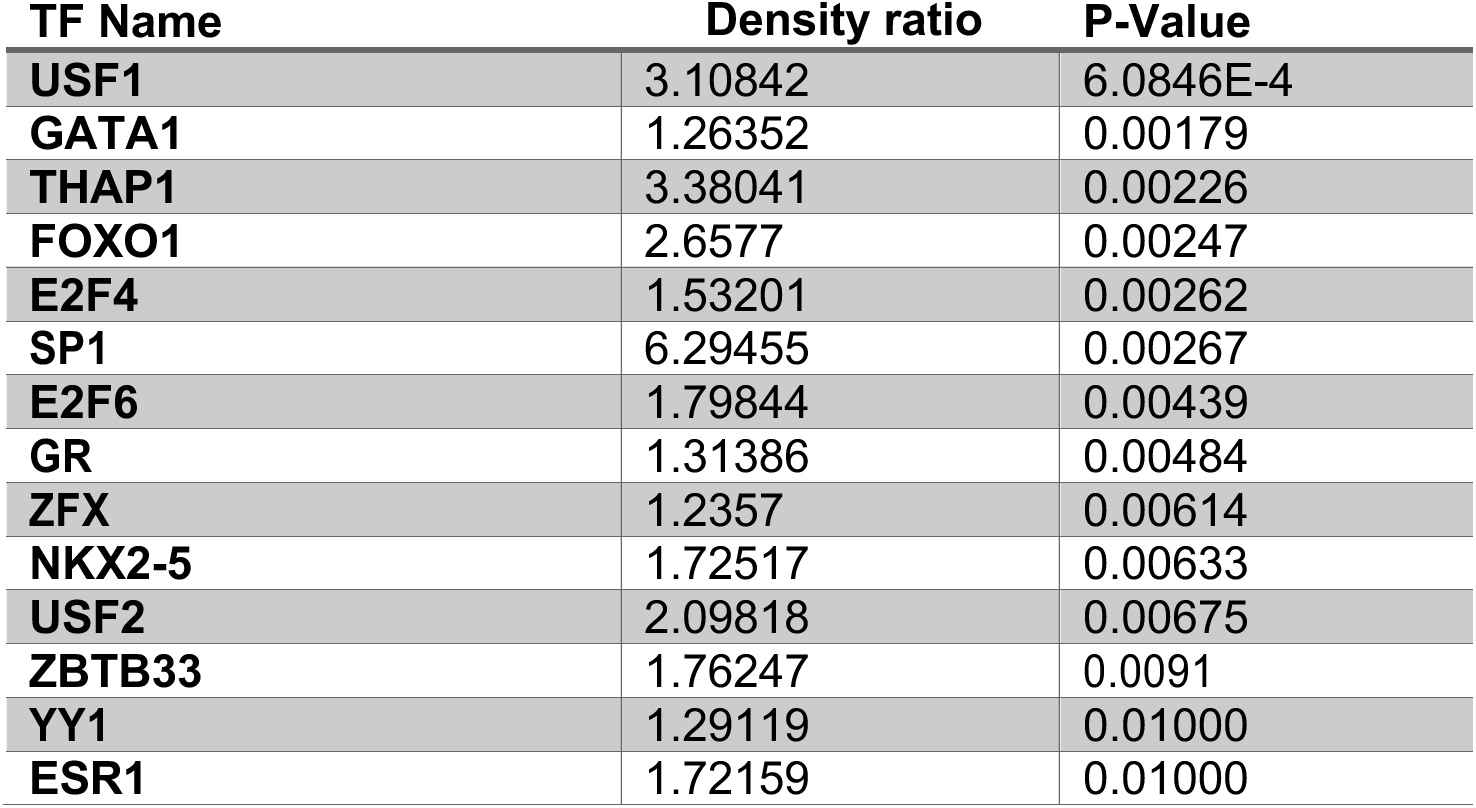
Transcription Motifs Enriched in DKO female livers compared to DKO males.

**Supplementary Table 4.**
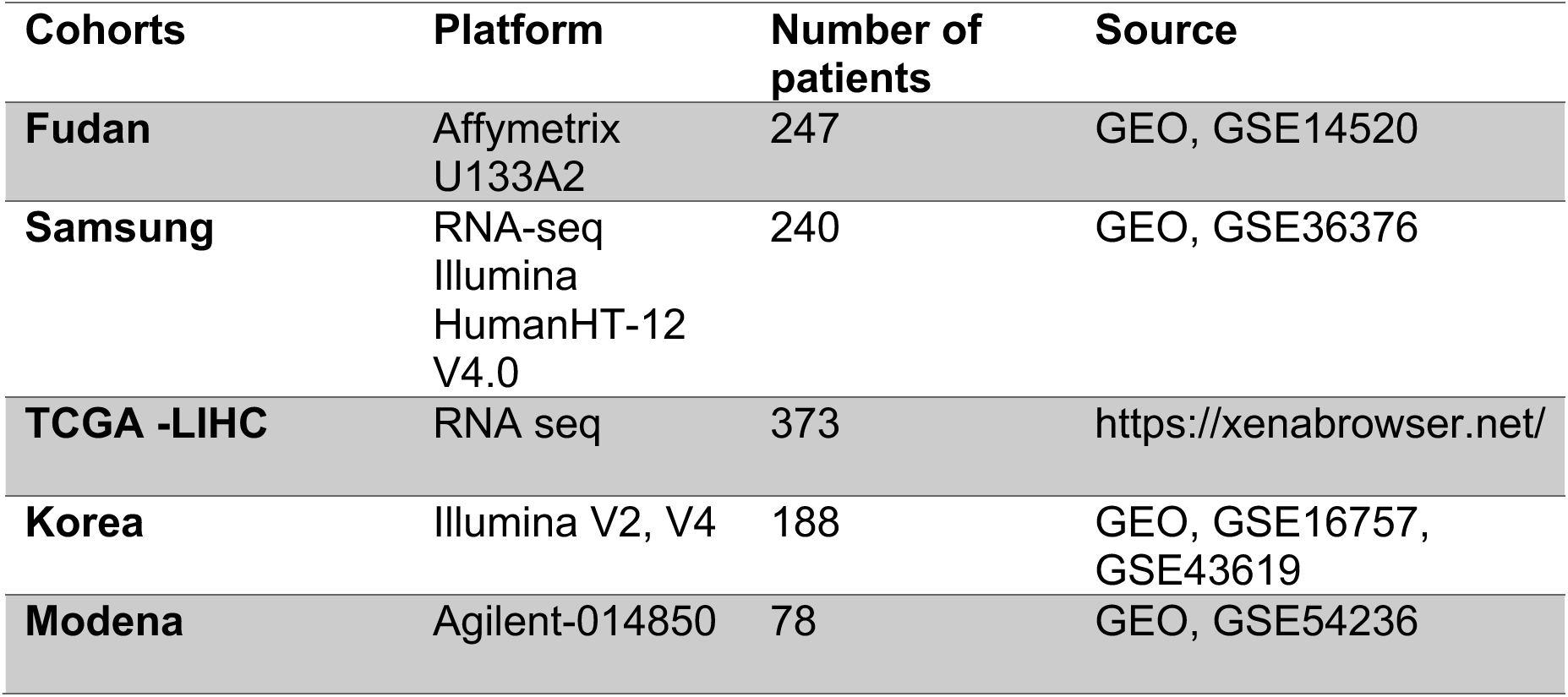
Summary of HCC gene expression data sets.

**Supplementary Table 5:**
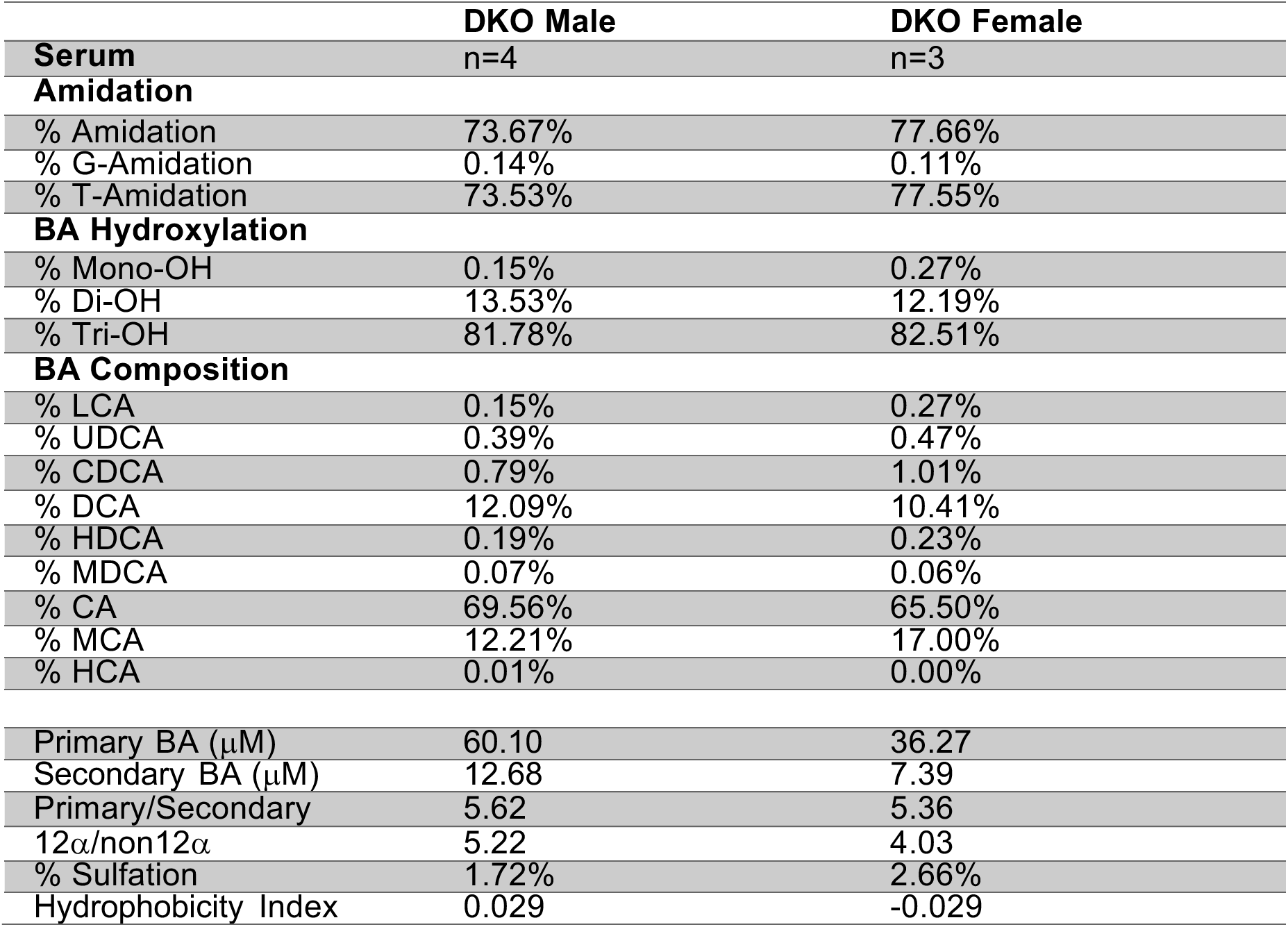
Serum BA composition in DKO mice.

| **Serum** | **DKO Male**<br>n=4 | **DKO Female**<br>n=3 |
| --- | --- | --- |
| **Amidation** |  |  |
| % Amidation | 73.67% | 77.66% |
| % G-Amidation | 0.14% | 0.11% |
| % T-Amidation | 73.53% | 77.55% |
| **BA Hydroxylation** |  |  |
| % Mono-OH | 0.15% | 0.27% |
| % Di-OH | 13.53% | 12.19% |
| % Tri-OH | 81.78% | 82.51% |
| **BA Composition** |  |  |
| % LCA | 0.15% | 0.27% |
| % UDCA | 0.39% | 0.47% |
| % CDCA | 0.79% | 1.01% |
| % DCA | 12.09% | 10.41% |
| % HDCA | 0.19% | 0.23% |
| % MDCA | 0.07% | 0.06% |
| % CA | 69.56% | 65.50% |
| % MCA | 12.21% | 17.00% |
| % HCA | 0.01% | 0.00% |
| Primary BA (μM) | 60.10 | 36.27 |
| Secondary BA (μM) | 12.68 | 7.39 |
| Primary/Secondary | 5.62 | 5.36 |
| 12α/non12α | 5.22 | 4.03 |
| % Sulfation | 1.72% | 2.66% |
| Hydrophobicity Index | 0.029 | -0.029 |

**Supplementary Table 6.**
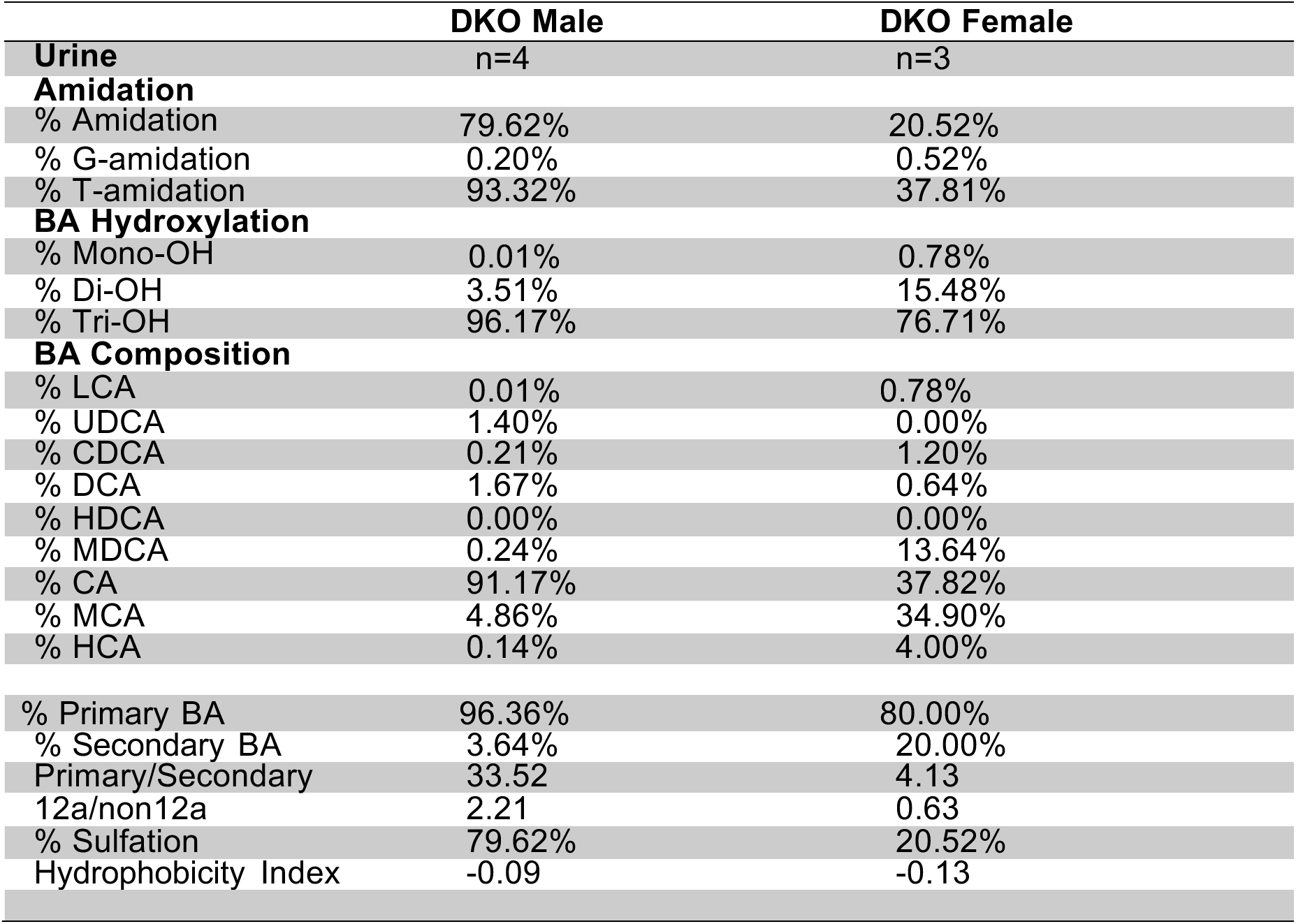
Urine BA composition in DKO mice.

**Supplementary Table 7.**
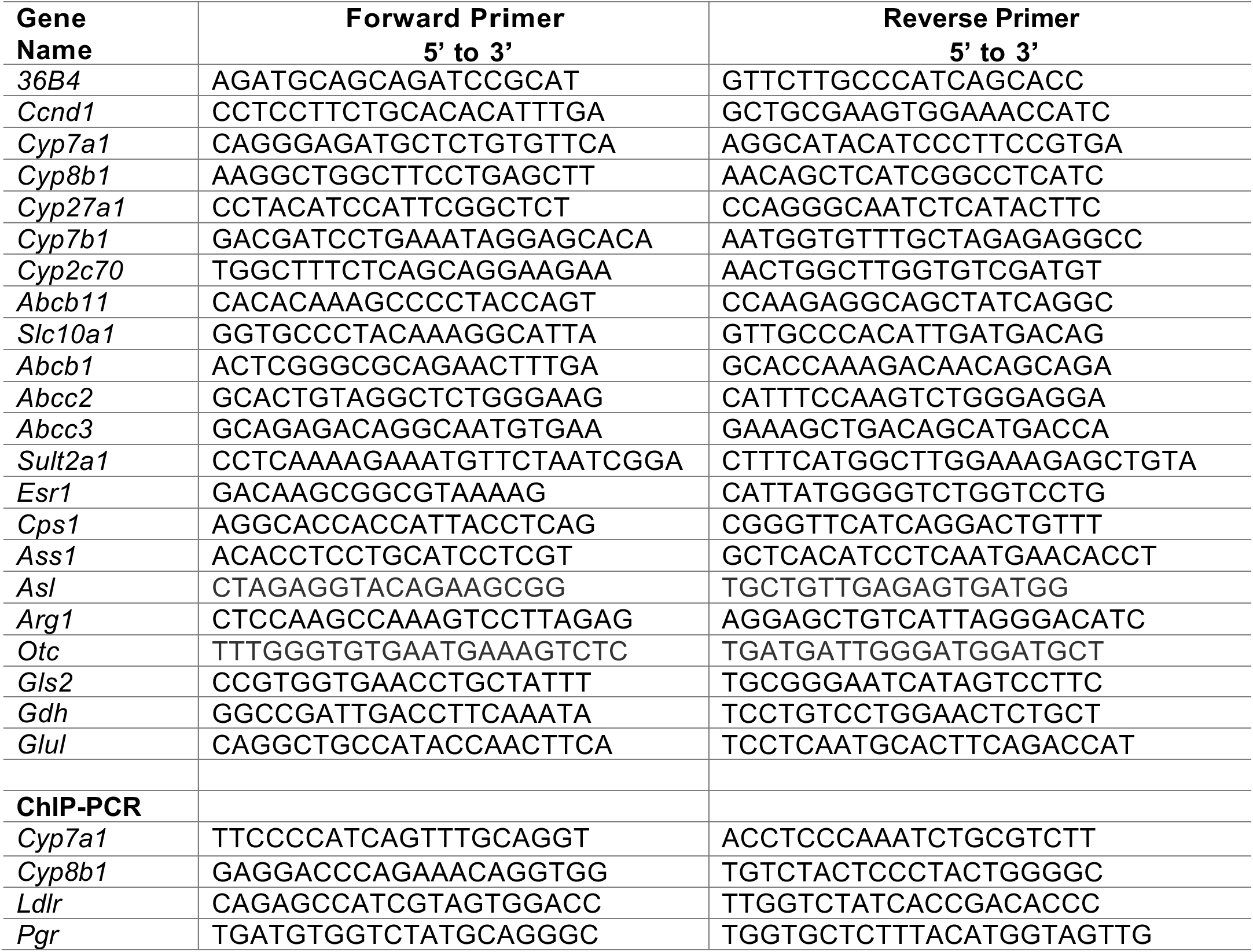
Primer sequences used.

