## Supplementary Tables for "Sex differences in bile acid homeostasis and excretion underlie the disparity in liver cancer incidence between males and females"

**Supplementary Table 1:**

**List of genes in different DKO gene signature category used for analysis**
**See Excel sheet**

**Supplementary Table 2. Transcription Motifs Enriched in DKO male livers compared to DKO females.**

| TF Name | Density ratio | P-Value |
|---|---|---|
| **PPARγ** | 1.30126 | 6.3527E-11 |
| **TCF3** | 1.33458 | 7.2023E-11 |
| **TCF4** | 1.27146 | 5.5608E-9 |
| **p73** | 1.30219 | 1.1841E-5 |
| **LRH1** | 1.1917 | 2.3061E-5 |
| **AR** | 1.3065 | 2.7733E-5 |
| **FOXA2** | 1.21861 | 6.526E-4 |
| **MEF2A** | 1.35309 | 0.00153 |
| **MEF2C** | 1.34771 | 0.0016 |
| **TBR2 (EOMES)** | 1.15298 | 0.00168 |
| **TR4** | 1.45287 | 0.00195 |
| **GATA3** | 1.95407 | 0.00351 |
| **c-JUN** | 1.20405 | 0.00491 |
| **STAT1** | 1.17336 | 0.00505 |
| **RXRa** | 1.42981 | 0.00512 |
| **STAT5a** | 1.19101 | 0.00557 |
| **FOXA1** | 1.53342 | 0.00674 |
| **STAT4** | 1.42981 | 0.00707 |
| **NRF2** | 3.06387 | 0.00959 |
| **KLF4** | 1.08893 | 0.01000 |

**Supplementary Table 3. Transcription Motifs Enriched in DKO female livers compared to DKO males.**

| TF Name | Density ratio | P-Value |
|---|---|---|
| **USF1** | 3.10842 | 6.0846E-4 |
| **GATA1** | 1.26352 | 0.00179 |
| **THAP1** | 3.38041 | 0.00226 |
| **FOXO1** | 2.6577 | 0.00247 |
| **E2F4** | 1.53201 | 0.00262 |
| **SP1** | 6.29455 | 0.00267 |
| **E2F6** | 1.79844 | 0.00439 |
| **GR** | 1.31386 | 0.00484 |
| **ZFX** | 1.2357 | 0.00614 |
| **NKX2-5** | 1.72517 | 0.00633 |
| **USF2** | 2.09818 | 0.00675 |
| **ZBTB33** | 1.76247 | 0.0091 |
| **YY1** | 1.29119 | 0.01000 |
| **ESR1** | 1.72159 | 0.01000 |

**Supplementary Table 4. Summary of HCC gene expression data sets**

| Cohorts | Platform | Number of patients | Source |
|---|---|---|---|
| **Fudan** | Affymetrix U133A2 | 247 | GEO, GSE14520 |
| **Samsung** | RNA-seq Illumina HumanHT-12 V4.0 | 240 | GEO, GSE36376 |
| **TCGA -LIHC** | RNA seq | 373 | https://xenabrowser.net/ |
| **Korea** | Illumina V2, V4 | 188 | GEO, GSE16757, GSE43619 |
| **Modena** | Agilent-014850 | 78 | GEO, GSE54236 |

**Supplementary Table 5: Serum BA composition in DKO mice.**

| | DKO Male | DKO Female |
|---|---|---|
| **Serum** | n=4 | n=3 |
| **Amidation** | | |
| % Amidation | 73.67% | 77.66% |
| % G-Amidation | 0.14% | 0.11% |
| % T-Amidation | 73.53% | 77.55% |
| **BA Hydroxylation** | | |
| % Mono-OH | 0.15% | 0.27% |
| % Di-OH | 13.53% | 12.19% |
| % Tri-OH | 81.78% | 82.51% |
| **BA Composition** | | |
| % LCA | 0.15% | 0.27% |
| % UDCA | 0.39% | 0.47% |
| % CDCA | 0.79% | 1.01% |
| % DCA | 12.09% | 10.41% |
| % HDCA | 0.19% | 0.23% |
| % MDCA | 0.07% | 0.06% |
| % CA | 69.56% | 65.50% |
| % MCA | 12.21% | 17.00% |
| % HCA | 0.01% | 0.00% |
| Primary BA ($\mu$M) | 60.10 | 36.27 |
| Secondary BA ($\mu$M) | 12.68 | 7.39 |
| Primary/Secondary | 5.62 | 5.36 |
| 12$\alpha$/non12$\alpha$ | 5.22 | 4.03 |
| % Sulfation | 1.72% | 2.66% |
| Hydrophobicity Index | 0.029 | -0.029 |

**Supplementary Table 6. Urine BA composition in DKO mice.**

| | DKO Male | DKO Female |
|---|---|---|
| **Urine** | n=4 | n=3 |
| **Amidation** | | |
| % Amidation | 79.62% | 20.52% |
| % G-amidation | 0.20% | 0.52% |
| % T-amidation | 93.32% | 37.81% |
| **BA Hydroxylation** | | |
| % Mono-OH | 0.01% | 0.78% |
| % Di-OH | 3.51% | 15.48% |
| % Tri-OH | 96.17% | 76.71% |
| **BA Composition** | | |
| % LCA | 0.01% | 0.78% |
| % UDCA | 1.40% | 0.00% |
| % CDCA | 0.21% | 1.20% |
| % DCA | 1.67% | 0.64% |
| % HDCA | 0.00% | 0.00% |
| % MDCA | 0.24% | 13.64% |
| % CA | 91.17% | 37.82% |
| % MCA | 4.86% | 34.90% |
| % HCA | 0.14% | 4.00% |
| % Primary BA | 96.36% | 80.00% |
| % Secondary BA | 3.64% | 20.00% |
| Primary/Secondary | 33.52 | 4.13 |
| 12a/non12a | 2.21 | 0.63 |
| % Sulfation | 79.62% | 20.52% |
| Hydrophobicity Index | -0.09 | -0.13 |

**Supplementary Table 7. Primer sequences used**

| Gene Name | Forward Primer 5' to 3' | Reverse Primer 5' to 3' |
|---|---|---|
| *36B4* | AGATGCAGCAGATCCGCAT | GTTCTTGCCCATCAGCACC |
| *Ccnd1* | CCTCCTTCTGCACACATTTGA | GCTGCGAAGTGGAAACCATC |
| *Cyp7a1* | CAGGGAGATGCTCTGTGTTCA | AGGCATACATCCCTTCCGTGA |
| *Cyp8b1* | AAGGCTGGCTTCCTGAGCTT | AACAGCTCATCGGCCTCATC |
| *Cyp27a1* | CCTACATCCATTCGGCTCT | CCAGGGCAATCTCATACTTC |
| *Cyp7b1* | GACGATCCTGAAATAGGAGCACA | AATGGTGTTTGCTAGAGAGGCC |
| *Cyp2c70* | TGGCTTTCTCAGCAGGAAGAA | AACTGGCTTGGTGTCGATGT |
| *Abcb11* | CACACAAAGCCCCTACCAGT | CCAAGAGGCAGCTATCAGGC |
| *Slc10a1* | GGTGCCCTACAAAGGCATTA | GTTGCCCACATTGATGACAG |
| *Abcb1* | ACTCGGGCGCAGAACTTTGA | GCACCAAAGACAACAGCAGA |
| *Abcc2* | GCACTGTAGGCTCTGGGAAG | CATTTCCAAGTCTGGGAGGA |
| *Abcc3* | GCAGAGACAGGCAATGTGAA | GAAAGCTGACAGCATGACCA |
| *Sult2a1* | CCTCAAAAGAAATGTTCTAATCGGA | CTTTCATGGCTTGGAAAGAGCTGTA |
| *Esr1* | GACAAGCGGCGTAAAAG | CATTATGGGGTCTGGTCCTG |
| *Cps1* | AGGCACCACCATTACCTCAG | CGGGTTCATCAGGACTGTTT |
| *Ass1* | ACACCTCCTGCATCCTCGT | GCTCACATCCTCAATGAACACCT |
| *Asl* | CTAGAGGTACAGAAGCGG | TGCTGTTGAGAGTGATGG |
| *Arg1* | CTCCAAGCCAAAGTCCTTAGAG | AGGAGCTGTCATTAGGGACATC |
| *Otc* | TTTGGGTGTGAATGAAAGTCTC | TGATGATTGGGATGGATGCT |
| *Gls2* | CCGTGGTGAACCTGCTATTT | TGCGGGAATCATAGTCCTTC |
| *Gdh* | GGCCGATTGACCTTCAAATA | TCCTGTCCTGGAACTCTGCT |
| *Glul* | CAGGCTGCCATACCAACTTCA | TCCTCAATGCACTTCAGACCAT |
| **ChIP-PCR** | | |
| *Cyp7a1* | TTCCCCATCAGTTTGCAGGT | ACCTCCCAAATCTGCGTCTT |
| *Cyp8b1* | GAGGACCCAGAAACAGGTGG | TGTCTACTCCCTACTGGGGC |
| *Ldlr* | CAGAGCCATCGTAGTGGACC | TTGGTCTATCACCGACACCC |
| *Pgr* | TGATGTGGTCTATGCAGGGC | TGGTGCTCTTTACATGGTAGTTG |
